# Data-driven polymer modeling reveals how scale-dependent active fluctuations shape chromatin organization

**DOI:** 10.64898/2026.04.27.721044

**Authors:** PP Adwaith, Sangram Kadam, Siddharth Pankar, Dimple Notani, Ranjith Padinhateeri

## Abstract

The nature of the chromatin polymer and its properties are tightly coupled to its function. By analyzing recent microscopy data, we show that chromatin segments at ∼ 10–100 kb scales exhibit anomalously broad bond-length fluctuations and acute local angles beyond the scope of existing polymer models. Using polymer simulations at nucleosome resolution and systematic coarse-graining, we show that chromatin at these scales requires a non-equilibrium description. We develop a data-constrained non-equilibrium model that explains these anomalous fluctuations, in which coarse-grained beads experience extensile, angular, and stochastic active forces. Our work provides an experimentally guided framework for incorporating active processes and suggests that the nature of activity is scale-dependent. The model quantitatively reproduces three-dimensional distance distributions across scales and enables inference of effective elastic and active parameters, providing a unified framework for active chromatin simulations and the study of its 3D organization.

## Introduction

The polymeric, sequential arrangement of bases in DNA encodes the genetic code. However, several cellular processes, including transcription, DNA replication, and repair, are regulated by the folded polymer known as chromatin. Chromatin exhibits a hierarchical organization, with the basic level arising from the wrapping of DNA around histone proteins to form a continuous chromatin fiber composed of nucleosomes. This fiber is further folded into multiple layers of loops and domains across scales, giving rise to a functional three-dimensional (3D) genome organization [1, 2]. This spatial organization is highly dynamic; its establishment and maintenance require a multi-layered workforce of ATP-dependent molecular machines operating across scales [2, 3].

A widely used experimental approach for probing the 3D organization of chromatin is the high-throughput chromatin conformation capture (Hi-C) method [4]. Hi-C provides population-averaged contact maps of chromatin organization, typically at resolutions on the order of a few kilobases [5]. Based on these data, chromatin is commonly described as a polymer organized into topologically associating domains (TADs), each spanning several hundred kilobases, and into compartments arising from the preferential intermixing of TADs with similar epigenetic states [5–7].

High-resolution microscopy provides a more direct view of chromatin organization, albeit with limitations [8– 13]. Recently developed oligopaint-based approaches allow tracing of chromatin polymer configurations in fixed cells [8, 14]. Using this method, Bintu et al. [8] reconstructed three-dimensional contours of a few Mb region of human chromosome 21 at 30 kb resolution, while Su et al. [14] reported configurations of larger chromatin regions at 50 kb resolution. Recently, Murphy and Boettiger obtained 3D configurations of a polycomb-bound heterochromatin region in mouse embryonic stem cells [15]. These measurements yield polymer configurations for thousands of individual cells, enabling comparisons between single-cell chromatin structures and population-averaged descriptions. While tracing approaches are limited to fixed cells, live-cell microscopy enables measurements of chromatin dynamics, i.e., time-dependent changes in chromatin organization [9, 10, 12, 16, 17, 32]. Such experiments typically track the relative motion of a small number of selected chromosomal loci. They show that chromatin loci undergo continuous motion over time, and how different proteins (e.g., CTCF and cohesin) influence these dynamics.

At smaller length scales, Micro-C experiments suggest that chromatin is organized into TAD-like domains and sub-TADs down to ∼10 kb [18–20]. Other studies indicate that nucleosomes form small clusters or clutches at the kilobase scale [20–23]. Contact probabilities among looped segments are relatively low (∼0.1), indicating that these structures are highly dynamic [3, 24]. Chromatin segments experience multiple sources of active forces, including continuous nucleosome sliding, disassembly and re-assembly [25, 26], transcriptional activity [27–32], loop extrusion [33–35], and ongoing fluctuations in histone modification states [36–38] all of which can lead to fluctuations in local chromatin structure.

Chromatin is commonly modeled as a bead–spring polymer with heterogeneous intra-chromatin interactions that capture looping between distant genomic loci [39–45]. Over the past decade, significant effort has focused on characterizing these interactions, either through effective attractive potentials between chromatin segments or via explicit protein-mediated bridging mechanisms [40, 42, 44–50]. More recently, models incorporating loop extrusion and interactions mediated by cohesin and CTCF have provided an additional mechanistic framework [27, 33, 34, 51–53]. Despite these advances, a fundamental aspect of chromatin polymer models remains poorly constrained: the elastic properties of the in vivo polymer itself. Nearly all existing models represent connectivity between neighboring coarsegrained chromatin segments using linear/non-linear springs, yet the associated elastic parameters are typically chosen in an ad hoc manner, without quantitative experimental grounding. Similarly, the in vivo bending fluctuations at local chromatin scales remain largely unexplored. While nucleosome-scale organization, transient looping, and local contact probabilities are expected to influence bending elasticity, these effects have not been systematically incorporated into current models.

This gap is further significant, given that chromatin is continuously driven by ATP-dependent processes [3, 30, 45, 54, 55]. As a result, the effective stretching and bending elastic properties of chromatin are emergent quantities shaped by the interplay between active processes and chromatin interactions. Determining these properties, and developing quantitative frameworks to infer them from experimental data, remains an important open problem. Accurate description of chromatin elasticity is essential for quantitatively predicting three-dimensional distances between functional genomic elements, such as enhancer–promoter pairs and distal genes, and hence for understanding the physical basis of gene regulation. Recent high-resolution microscopy experiments further challenge existing paradigms by revealing unexpectedly large fluctuations in the spatial extent of chromatin segments (∼ 10–100 kb), far exceeding predictions from standard equilibrium polymer models [8, 14, 15]. These observations point to a fundamental limitation of current descriptions and highlight the need for a novel approach.

Here, we address this gap by systematically analyzing recent microscopy data and developing a coarse-grained, non-equilibrium polymer model for chromatin. We demonstrate that incorporating activity at the level of chromatin segments naturally explains the observed large fluctuations. Importantly, our framework enables the quantitative inference of effective elastic parameters, as well as the strength and characteristics of active processes, consistent with experimental data. Our results further reveal that chromatin activity is inherently scale-dependent, with directed forces at short length scales giving rise to effectively random behavior at larger scales. Our model provides a unified and physically consistent description across scales—from nucleosome-level organization to highly coarse-grained chromatin segments—and quantitatively reproduces not only mean three-dimensional distances but also the full distribution of distances between genomic loci, offering a stringent test of model validity. Finally, we propose plausible microscopic mechanisms, such as nucleosome disassembly and protein-mediated state switching, that can give rise to the observed non-equilibrium fluctuations. Together, these results establish that chromatin mechanics must be understood within an active, non-equilibrium framework and enable quantitative extraction of its effective material properties.

## Results

### Estimating human chromatin polymer parameters and predicting 3D distances

One of the major challenges in chromatin simulations is the lack of well-defined polymer parameters for human chromatin. To perform coarse-grained chromatin polymer simulations at the scale of a few TADs (megabases) or larger, one requires parameters such as stretching elasticity, bending elasticity, and intra-chromatin interaction potentials. Since the nucleosome structure and physical dimensions are known at the 100–200 bp scale, one approach is to build a bottom-up model starting from nucleosome resolution and coarse-grain the polymer to obtain properties at larger scales [56].

We started with nucleosome-resolution (200bp) Micro-C data for human chromosome [19] and generated an ensemble of 3D chromatin configurations using our MicroC-based Chromatin Model (MCCM) [56]. The method involves implementing intra-chromatin contacts by connecting selected pairs of nucleosomes via harmonic springs with probability *P*_*ij*_ consistent with the contact map data, and physical proximity (see Fig. 1A, Methods MCCM, and Supplementary Information (SI) Text S1 A). Then the system — chromatin at nucleosome resolution — is “equilibrated” using Langevin dynamics simulations. The resulting contact map from simulations matches well with the experimental contact map (Fig. 1B). This agreement is quantified using the stratum-adjusted correlation coefficient (SCC = 0.93) (see SI Text S1 I) [57]. From these simulated configurations we can predict mean 3D distance between any two 200bp segments (Fig. 1C-D).

**FIG. 1:**
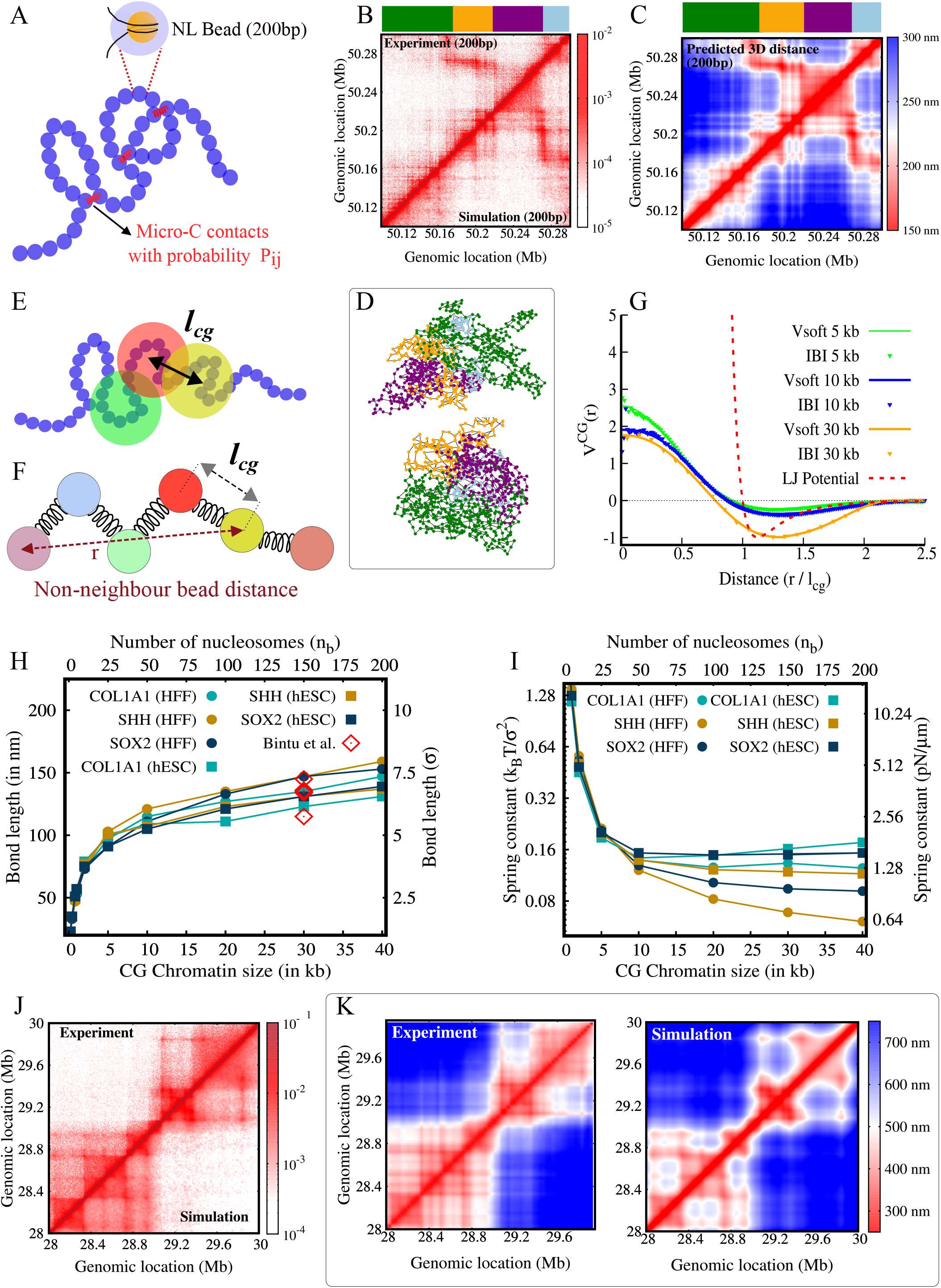
Computing coarse-grained (CG) polymer parameters for human chromatin: (A) Schematic of chromatin represented as a polymer, where each bead corresponds to a nucleosome+linker (NL, ∼200bp). Beads interact via intra-chain springs with probabilities *P*_*ij*_ derived from Micro-C data [19]. Ensembles of 3D structures are generated using Langevin dynamics simulations, from which pairwise 3D distances are computed. (B) Using the human Micro-C contact map (HFF, COL1A1 gene region, chr17: 50.1Mb-50.3Mb) [19], we generated an ensemble of 3D structures. The contact map obtained from simulations agrees well with the experimental map. (C) Predicted 3D distances between 200bp chromatin segments, with size of the 200bp region (*σ*) is taken as 20nm (see SI text S1 C). (D) Two representative polymer conformations from the simulations; colors correspond to the segments indicated in the strip above the contact and distance maps. (E) Schematic of the coarse-graining approach. Consecutive *n*_*b*_ nucleosome beads are grouped into a single CG bead. *l*_*cg*_ denotes the bond length between two neighboring CG beads. (F) Corresponding bead-spring polymer picture. CG beads are connected by springs; 3D distance between neighboring and non-neighboring beads (genomic segments) are represented by *l*_cg_ and *r* respectively. (G) Effective non-bonded soft potential (*V*_soft_) among beads for different CG sizes. Repulsive part is much softer compared to the Lennard–Jones (LJ) potential. (H) Most probable *l*_*cg*_ predicted from simulations as a function of coarse-graining size (*n*_*b*_) for multiple gene regions; the values are consistent with microscopy measurements (open diamonds) [8]. (I) The inverse of the *l*_cg_ variance is the effective spring constant (*K*_cg_) and is plotted as a function of coarse-graining size for multiple gene regions. Performed CG simulation of IMR90 chr21: 28–30 Mb region at 5kb resolution. The simulated (J) contact maps, and (K) 3D distances compare well with experiments.

We then systematically coarse-grained the nucleosome-resolution polymer by taking *n*_*b*_ consecutive nucleosomes as one big coarse-grained (CG) bead (see Fig. 1E-F, Methods) and generated a CG polymer. We define the bond-length of the CG polymer (*l*_cg_) as the 3D distance between two neighboring CG beads and compute it for any gene region (Fig. 1E-F). The most probable *l*_cg_ as a function of coarse-graining size (*n*_*b*_) for different gene regions are shown in Fig. 1H. This is the effective size of the region. When 30kb of chromatin (*n*_*b*_ = 150 nucleosomes) is coarse-grained as one bead, the bond length is in the range of 110-150 nm and this matches well with the 30kb inter-segmental distance obtained from super-resolution microscopy experiments in different human cell lines [8]. From the ensemble of configurations simulated, we compute the distribution of the bond length *P* (*l*_cg_) [56]. The inverse of the variance of the distribution gives us a measure of effective spring constant of the CG chromatin polymer, *K*_cg_ (Fig. 1I and SI Text S1 I). Starting from Micro-C data, our procedure predicts typical CG bead size (*l*_cg_) and spring constant (*K*_cg_) for human chromatin at any desired coarse-graining scale.

If one has to do a CG human chromatin polymer simulation, what should be the intra-chromatin interaction between the CG polymers? To compute this, we employed the iterative Boltzmann inversions method [56]; that is, we started with a CG chromatin polymer with an arbitrary intra-chromatin interaction potential and iteratively evolved the potential such that the non-bonded distance distribution among the CG beads is comparable to what is obtained from the nucleosome-resolution Micro-C derived polymer (see SI Text S1 B). This gives us a potential energy function as shown in Fig. 1G, which is highly soft compared to the typical Lennard-Jones (LJ) potential. The softness in the repulsive nature of the potential, unlike LJ, indicates that long chromatin polymer segments can mix with each other [58].

The coarse-graining procedure employed above gives us a good estimate of essential parameters — bead size, spring constant, intra-chromtin potential — to simulate human chromatin at large length-scale (Figs. 1G, H, I, and SI Table S2). Using these parameters we simulated a coarse-grained chromatin polymer representing the region Chr 21: 28-30Mb, IMR90, so that our predictions can be compared with available microscopy data [8]. The details of the simulation are as follows: We generated a CG chromatin bead-spring polymer with each bead as 5kb with relevant CG parameters as mentioned above. To account for specific intra-chromatin interactions, we inserted harmonic spring between specific segments based on 5kb Hi-C contact probability [5] (SI Text S1 A). The repulsive part of the intra-chromatin interaction was taken from the soft-potential above. This HiC-based Chromatin Simulation Method (HCCM, see Methods, and SI Text S1 A) gave us an ensemble of 3D polymer configurations for the entire 2Mb region. The contact map we obtained from simulations are comparable with the Hi-C contact map (SCC=0.98) (Fig. 1J). Our simulations predict the 3D distance between any two segments and this match well with the chromatin-tracing experimental data from Bintu et al. [8] (Fig. 1K). The correlation between the experimental data (in X-axis) and the predicted data from simulation (Y-axis) is evident in Fig. S2.

### Current polymer models cannot reproduce bond-length and bond-angle distributions observed in experiments

We analyzed existing single-cell chromatin tracing super-resolution microscopy data from refs. [8, 14] and computed the probability distribution of inter-segment distances, *P* (*l*_*cg*_), and the distribution of bending angles, *P* (*θ*_*cg*_) (Fig. 2). The experimental data provide the 3D positional coordinates of the centers of each 30 kb chromatin segment along the contour; these correspond to coarse-grained beads of size 30 kb in the polymer model. The distance distributions between adjacent 30 kb segments from experiments and our simulations are shown in Fig. 2A (bottom). The simulations are performed using the HCCM method described above (Methods and SI Text S1 A). Note that the most probable value of *l*_*cg*_ predicted from simulations is comparable to that measured in experiments. However, the experimental distance distribution exhibits a long tail that is absent in our simulation results. This tail suggests that the 30 kb chromatin polymer attains extended conformations with a small but finite probability. Such extended configurations have also been reported recently in ref. [15].

**FIG. 2:**
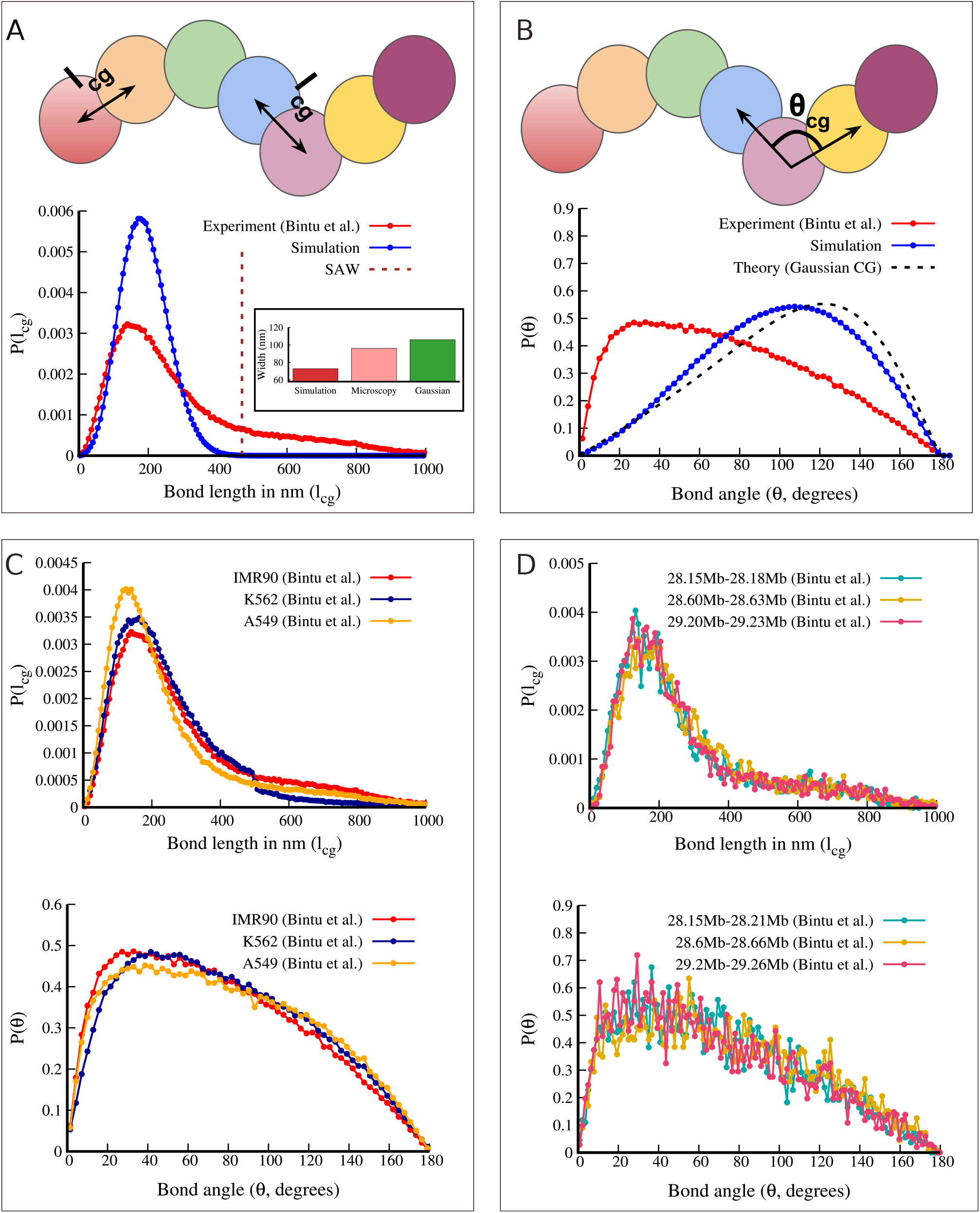
Current polymer models cannot reproduce bond-length and bond-angle distributions observed in microscopy: (A) Schematic representation of the bond length (*l*_cg_) (top). *P* (*l*_*cg*_) from our simulations compared with microscopy measurements from [8] for IMR90 chr21: 28–30 Mb (bottom). Although the most probable values match, there is a clear mismatch as experimental data shows a pronounced long tail. The dashed vertical line indicates the mean bond length of an equivalent random self-avoiding walk polymer, highlighting the extreme extensions observed. Inset: the width near the peak (a measure of fluctuations) shows that experimental configurations exhibit large fluctuations, comparable to a Gaussian (random) polymer. (B) Schematic representation of the bond angle (top) and the corresponding probability distributions P(*θ*_cg_) from simulations and experiments [8]. Theoretical curve for a CG Gaussian polymer [59] is shown as a guide. *P* (*l*_cg_) and *P* (*θ*_cg_) from different (C) cell lines, and (D) genomic locations [8] show a consistent pattern suggesting that this is general feature of chromatin segments.

To understand the nature of the extended configurations, we computed the mean extension of an equivalent random polymer generated via self-avoiding walk (SAW), and marked it as the dashed vertical line in Fig. 2A. This suggests that the observed extended configurations are as large as those of a SAW polymer (SI Text S1 A). This implies that, to attain such configurations, local monomers must collectively break all intra-chromatin interactions. From high-resolution contact map data, we know that a typical 30 kb chromatin segment contains several intra-chromatin bonds [5, 18–20, 56]. Breaking all these interactions simultaneously is improbable under equilibrium-like conditions and is likely to violate detailed balance.

These experimental data also provide information relevant for chromatin polymer models, particularly the nature of interactions between nearest-neighbor coarse-grained segments. Most coarse-grained chromatin models assume that neighboring segments can be represented as beads connected by a spring potential, typically harmonic or Finite Extensible Nonlinear Elastic (FENE). However, the experimental data indicate that the effective spring behavior is neither purely harmonic nor FENE. The FENE spring requires increasingly large forces to extend as it approaches its maximum extension. In contrast, the long tail in the chromatin data suggests that the 30kb region extends (or flows) under a very small net force (Fig. S3). Very near to the peak, the fluctuations are approximately harmonic and we quantify this by measuring the full width at 88% of the peak height (equivalent to the standard deviation for a Gaussian) (see SI Text S1 I), as shown in the inset of Fig. 2A. However, the overall fluctuation in the experimental data is significantly larger than that predicted by simulations incorporating Hi-C contacts and is comparable to that of an equivalent 30 kb Gaussian polymer (an ideal bead–spring polymer without non-bonded interactions) (SI Text S1 A). The long tail and high width quantify the anomalous behavior of the experimental data.

We then considered 3 consecutive 30kb segments (CG beads *i, i* + 1, *i* + 2) and computed the angle distribution between the beads — as shown in Fig. 2B the bending angle *θ*_*cg*_ subtended between the two vectors connecting the three beads. The experimentally observed angle distribution is different from the distribution predicted by our simulation. The experiments suggest that small angles are much more probable than expected from a basic theory [59]. The low angle suggests that segments within a few tens of kb away often loop, overlap and mix. Even for an ideal polymer (which allows self-overlap and has no excluded volume) or a LJ polymer with attractive interactions between all beads, the angle distribution peaks at values far from those observed experimentally [56, 59]. This suggests that, in vivo, certain folding events lie beyond the scope of equilibrium descriptions.

To examine whether the large distances and small angles observed here are specific to certain cell lines or genomic regions, we analyzed additional microscopy data from Bintu et al. [8] and Su et al. [14]. For the same genomic region (chr21: 28–30 Mb), *P* (*l*_cg_) and *P* (*θ*_cg_) show similar patterns across different cell lines (Fig. 2C). Across all cell lines, the bond-length distribution exhibits a similar long tail, indicating that the unusual extension is persistent (Fig. 2C, top panel). Similarly, the bending-angle distribution shows comparable preference for acute angles (Fig. 2C, bottom panel). Are these non-trivial distributions specific to particular genomic locations, or are they a general feature of chromatin? To understand this, we computed the distributions for all loci within the IMR90 chr21: 28–30 Mb region and found that this behavior is consistent across all loci (Fig. S4). Distributions for three randomly selected regions are shown as examples in Fig. 2D. The same behavior persists across experiments with different resolutions [14] (see Fig. S5). Moreover, a recent study by Murphy and Boettiger shows that such unusually extended configurations exist in both polycomb-dominated heterochromatic regions and non-polycomb regions [15].

### An active chromatin polymer model consistent with experimentally observed angle and distance distributions

Since coarse-graining is essential to simulate long (∼ 200 Mb) chromatin polymer and the full genome (∼ 3000 Mb), the standard approach is to represent chromatin as a linear chain of CG beads connected by springs. If one considers 30kb of chromatin as one bead, the two neighbouring beads are expected to have a distance (bond length) distribution similar to what is observed in experiments, as shown in Fig. 2A. However, to the best of our knowledge, no current bead-spring model can reproduce such a nearest-neighbor bond length distribution.

Based on the following observations, we argue that a non-equilibrium active model is the appropriate model to explain the experimental data: (i) The extension in the tail exceeds that of an equivalent random polymer or self-avoiding walk (SAW) polymer that has no attractive interactions (Fig. 2A). High-resolution Micro-C/Hi-C data indicate that a typical 30kb chromatin segment contains multiple loops and folds arising from attractive interactions. This implies that even attaining a SAW-like open configuration would require the collective breaking of all these interactions and it is not possible under thermal equilibrium, given the intra-chromatin interactions (see later sections). Furthermore, the angle distribution shows significant mixing/overlap of *i*^th^ and *i* + 2^nd^ beads, leading to an unusual bias toward small angles far away from what an “effective equilibrium” model can justify [56]. Together, these observations point to the action of non-equilibrium forces. (ii) At nucleosome-scale, non-equilibrium processes are ubiquitous. Chromatin is subject to pervasive ATP-dependent activity: nucleosomes are repositioned or disassembled by remodelers such as SWI/SNF, CHD, ISWI, and INO80 [60]. Transcriptional processes induce local configurational changes; continuous enzymatic activity associated with histone modifications, along with the flux of DNA-binding proteins, can dynamically reorganize chromatin. These processes can generate active forces and locally extended configurations, in principle.

Motivated by all these considerations, we hypothesize that each coarse-grained chromatin segment (here 30kb) experiences effective non-equilibrium extensile forces. In addition, guided by the experimental angle distribution, we assume that triplets of beads experience active bending forces that bias the angle distribution toward smaller values. We incorporate these extensile and angle-dependent active forces into our new Activity-Driven Chromatin Model (ADCM, see Methods) and simulate the polymer as described below.

We randomly choose any three consecutive beads in the polymer and apply either an extensile force of magnitude *F*_ext_ or an angle force *F*_ang_, randomly, for a duration Δ*τ* (Fig. 3A). The extensile force increases the separation between nearest neighbors, while the angle force bends the segment to bring the first and third beads closer, mimicking looping and overlap in the 30kb–90kb range (see ADCM in Methods). Our objective is to determine the magnitudes of these forces such that the experimentally observed length and angle distributions are reproduced. A constant active force applied continuously cannot generate the long tail (Fig. S6). The long tail indicates that large forces (and hence large extensions) are rare. Accordingly, we apply an extensile force for a duration inversely proportional to its magnitude, 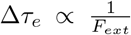, so that larger forces act only for short times. Similarly, the angle force is applied for a duration 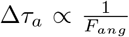. As described below (and in Methods), we simulate the polymer over a range of *F*_*ext*_ and *F*_*ang*_ values and identify the optimal set that reproduces the experimental distributions.

**FIG. 3:**
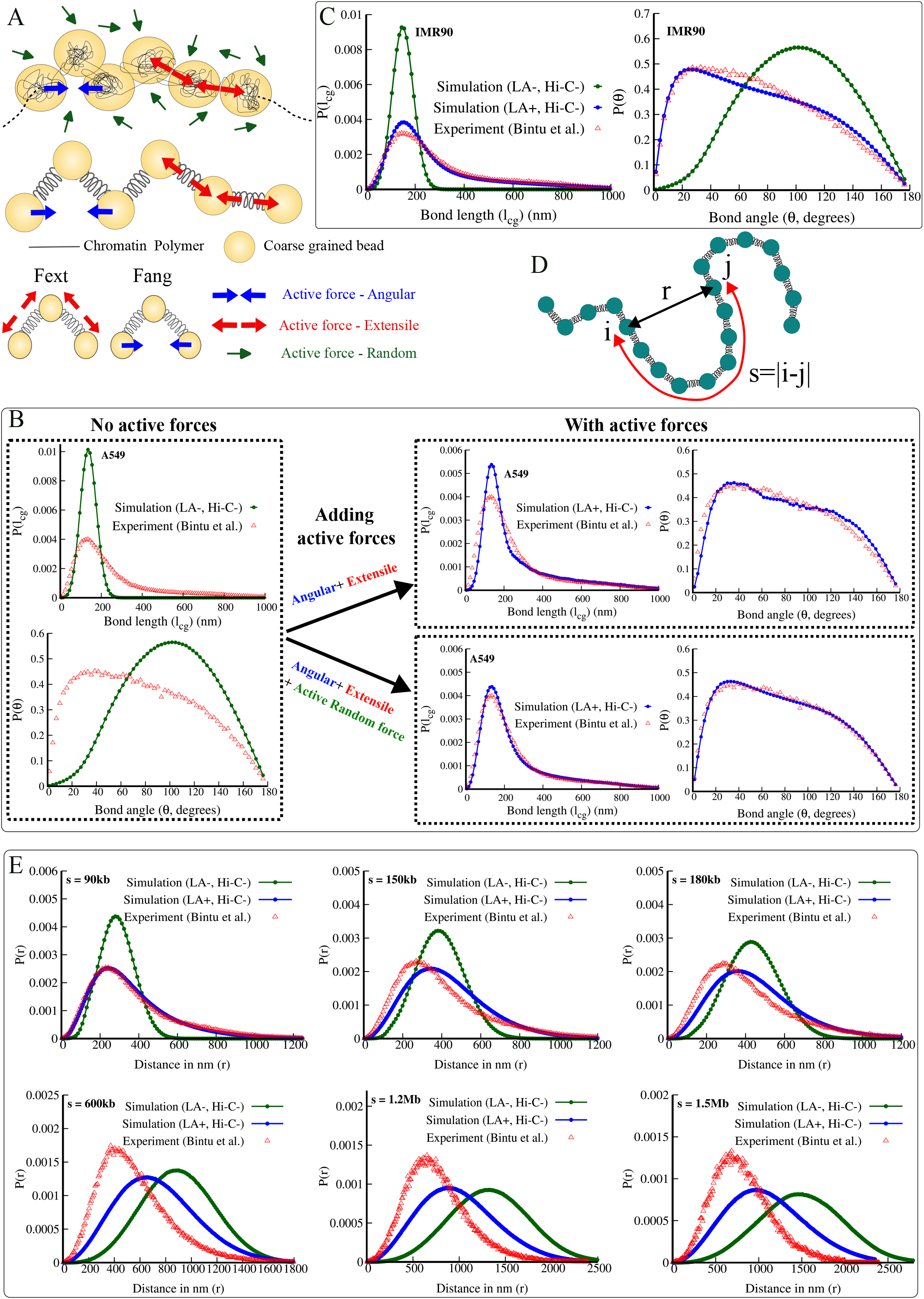
An active model consistent with experimentally observed bond-length and angle distributions: (A) Schematic of the proposed activity model for CG chromatin. Beads experience three types of active forces: extensile, angular, and random. Consecutive bead triplets are stochastically assigned either an extensile force (*F*_ext_, red arrows) or an angular force (*F*_ang_, blue arrows), each acting for a finite duration before being stochastically reassigned. In addition, a background random active force (green arrows) acts on the entire polymer throughout the simulation. (B) Application of extensile and angular active forces leads to a good fit to the angle distribution and the tail of the bond-length distribution; however, the width near the peak does not match (top right). All three active forces (extensile, angular, and random) are essential to obtain a good fit with experimental data (right bottom) [8]. (C) Comparison of the distributions obtained from simulations and microscopy for a different cell line (IMR90 [8]). (D) Schematic showing the genomic separation (s) between two segments *i* and *j* and the corresponding 3D distance (*r*). (E) 3D distance distributions between segments at fixed genomic separation (*s*). LA+*/*− denote simulations with/without local active forces, and Hi-C− indicates that long-range interactions observed in Hi-C (e.g., due to extrusion, CTCF, etc.) are not included.

To achieve this, we performed Brownian dynamics simulations of 250 beads polymer where the middle 67 beads represented a 2Mb region of chr 21 (28-30 Mb) such that each bead is 30kb. The rest of the beads are taken to avoid any small polymer size effects. Neighboring beads along the linear chromatin chain interact via a harmonic potential, with the CG spring constant taken from Fig. 1I. All non-bonded bead pairs interact through a self-avoiding soft potential, as described in Fig. 1G. To capture the long tail in the bond-length distribution and the skewness in the angle distribution, we incorporate the active extensile and angular forces as described above (see also Methods). Note that these active forces are local in nature, as they act only on three neighboring beads (referred to here as “local activity,” LA, acting in the range *<*100kb), in contrast to long-range processes such as loop extrusion.

We iteratively varied the active extensile and angular force parameters and obtained the best fit to the experimental data (Fig. 3B, top right). This yields *P* (*l*_cg_), where the tail of the distribution matches well with the experiment. However, the width near the peak is not well captured (width from simulation ≈50 nm, experiment ≈96 nm) indicating that chromatin segment size fluctuates more than predicted by this model. Given that non-equilibrium forces are pervasive at the nucleosome scale, a component of these forces can manifest as an effective random force at the 30kb scale, leading to enhanced fluctuations and a reduced effective spring constant. To account for this, in addition to the extensile and angular forces, we explicitly include active random forces in the model (Fig. 3A) quantified by an effective temperature-like parameter *T*_eff_ (see Methods). We therefore systematically varied three parameters—active random force, active extensile force, and active angular force—performing a 3D grid search over a broad range. The optimal parameters were obtained by minimizing the Jensen–Shannon divergence between simulated and experimental bond-length and angle distributions (see SI Text S1 D, SI Table S1). The resulting distributions are shown in Fig. 3B (right bottom panel). Note that, in this model so far, we do not include any long-range attractive interactions (e.g., cohesin-mediated loops) between distal genomic regions that are typically observed in Hi-C contact maps (Hi-C-). We also performed simulations for multiple other cell-lines as well with local activity (LA+) and without local activity (LA-) and found active parameters that match experiments (Fig. 3C and Fig. S7).

The optimal active force parameters for IMR90 lie in the following ranges: extensile forces *F*_*ext*_ are ≈ 1–11 pN. The angle forces are smaller, with *F*_*ang*_ in the range ≈ 0.37–3.73 pN. As noted above and in Methods, larger forces act with lower probability (shorter duration), while smaller forces act with higher probability (longer duration). The active random force values are such that *T*_eff_ */T* = 15.5. These values may not represent the action of individual molecules but rather the net effective forces acting over a coarse-grained 30kb segment. At smaller scales—such as the nucleosome or kilobase scale—forces may be larger but act in different directions, partially canceling and resulting in the effective magnitudes reported here. Since we do not have any attractive interactions among distal (*>* 90 kb) genomic regions in these simulations (HiC-), the results presented here demonstrate solely the effect of non-equilibrium active forces acting locally, in contrast to long-range interactions (e.g., extrusion or other mechanisms) that generate contacts observed in Hi-C maps. The effect of explicit non-local forces will be described in the next section.

Although the active forces act locally, we ask whether they influence distances between distal beads (|*i* − *j*| *>* 2) by propagating along the polymer network. We computed 3D distances (*r*) among two different beads (genomic segments *i, j*) that are separated by genomic distances, s (see Fig. 3D) ranging from 90kb to 1.5 Mb, and their distributions are plotted in Fig. 3E. We first compare simulations with local activity (LA+, blue) and without activity (LA-, green). Across all length scales, local activity alters the distributions. At short to intermediate scales (90kb, 150kb, and 180kb), pronounced long tails are observed in the LA+ case (blue curves). At larger scales (≈ 1 Mb), the distributions become more symmetric, suggesting that active extensile forces acting at 10–100 kb do not lead to directed extensile forces at megabase scales. Instead, directed active forces at short scales effectively cancel, giving rise to random-like fluctuations at larger scales. This indicates that chromatin organization at different length scales may be governed by distinct types of active forces. Next, we compare the LA+ simulation curves (blue) with experimental data (red) from Bintu et al. [8]. At 90kb (*i*–(*i* + 3) beads) and 150kb (*i*–(*i* + 5) beads), local activity alone brings the simulated distributions closer to the experimental ones. However, the simulations lack intra-chromatin attractive interactions between distant segments (e.g., mediated by CTCF, Cohesin or other inter-TAD/intra-compartment interactions), which give rise to Hi-C contacts. The absence of these interactions (Hi-C-) leads to less compact distributions in the simulations (blue) compared to experiments (red). Overall, we show that local activity alone reproduces the long tails and introduces partial compaction. However, additional long-range interactions are required to accurately capture chromatin organization at larger genomic distances.

### The chromatin model with activity and intra-chromatin interactions reproduces the full 3D distance distributions and fluctuations

In this section, we go beyond local activity and introduce interactions among distant chromatin segments (beyond *i*–(*i* + 2) beads) based on Hi-C data. To update Activity-Driven Chromatin Model (ADCM, Methods) with Hi-C-informed interactions, we select prominent contacts from the corresponding Hi-C contact maps [5] and introduce probabilistic harmonic interactions between the corresponding bead pairs (Fig. 4A; Methods). These attractive interactions often arise from underlying active processes (e.g., loop extrusion) acting between distant segments (see Fig. S8). We do Brownian Dynamics simulation of the system incorporating both local activity and HiC-informed attractive interactions (see Methods). The results are shown in Fig. 4B-F. Fig. 4B shows the bond-length and bond-angle distributions with and without local activity (LA+/-) in the presence of HiC-informed interactions (Hi-C+). Even in the presence of HiC-informed interactions, local activity (LA+) is necessary for formation of the long tail and peak at very small angles. In Fig. 4C, we compare contact probabilities between experiment and simulation for the 2Mb region; the agreement is very good (SCC=0.95). Our model further predicts the 3D distances (*r*) between any two segments, which also match well with experiments (Fig. 4D). A scatter plot of experimental versus simulated distances (Fig. 4E) shows strong correlation along the *X* = *Y* line. In Fig. 4F, we compare the distance distributions between distal genomic segments from simulations (blue) and experiments (red), demonstrating that not only the mean but the entire distribution can be predicted. This also highlights the role of activity. Even in the presence of HiC-informed interactions, the absence of activity (LA-, HiC+) leads to noticeable deviations from experiments (green vs red curve), indicating that the activity is essential for reproducing the full distribution. Comparing distributions between Figs. 3E and 4E, it is clear that for short genomic distances (e.g., s = 180*kb*), the activity produces the long tail in the distributions (extended configurations) while HiC-based interactions compact the polymer. However for distal regions (*>*Mb) both the forces compact chromatin as the local active forces are averaging out to nullify any extensile effects.

**FIG. 4:**
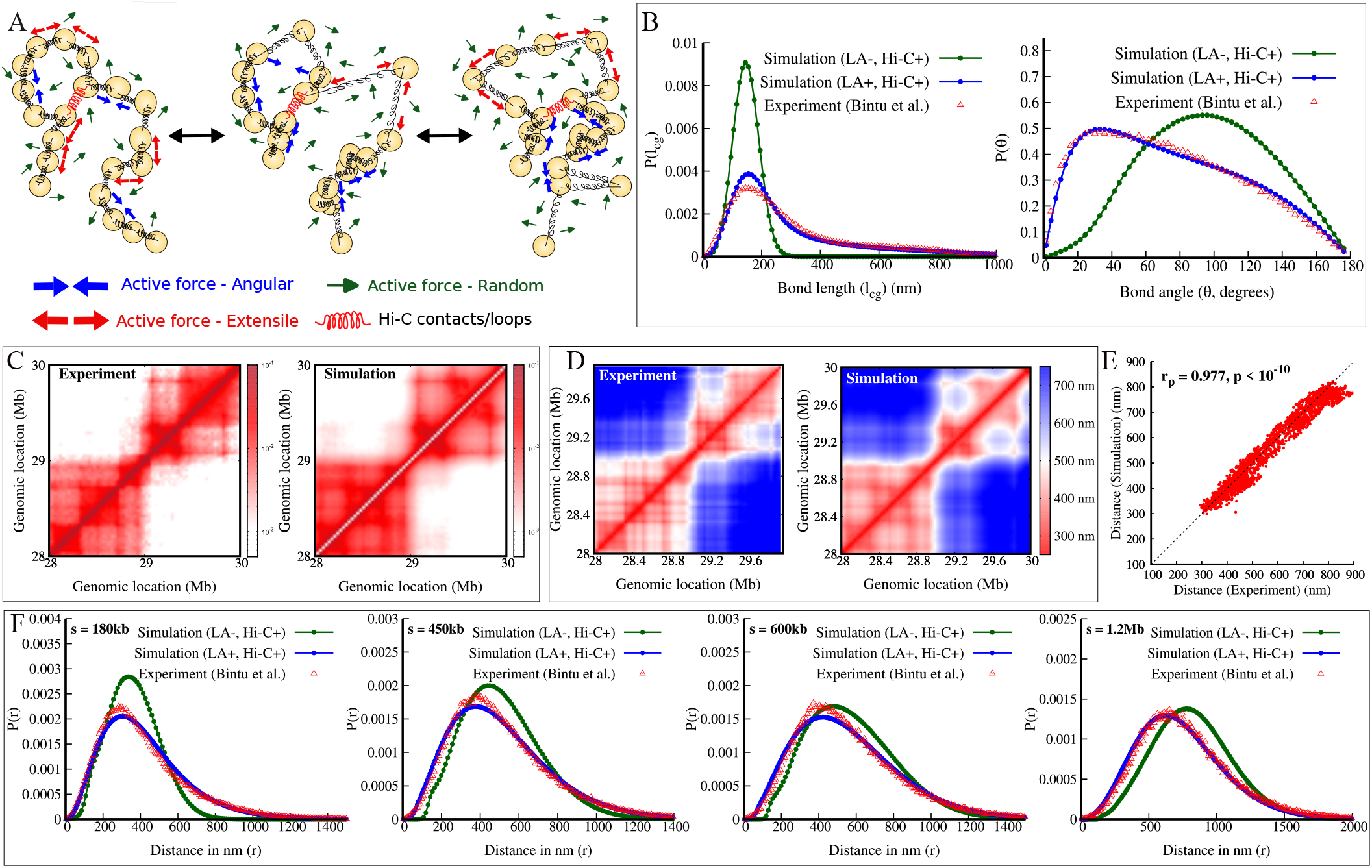
Model with activity and intra-chromatin interactions reproduces the full 3D distance distributions and local fluctuations: (A) Schematic of the coarse-grained polymer with active forces and HiC–informed interactions (red springs). (B) When simulations are performed with local active forces (LA+) and HiC–informed interactions (HiC+), the distributions from simulations match the experiments. In contrast, simulations without local active forces (LA-) show deviations. Comparison of the (C) contact maps and (D) mean 3D distances obtained from the active-model simulations and experiments. (E) Predicted 3D distances from simulations correlate well with the corresponding microscopy measurements. (F) 3D distance distributions between segments with large genomic separation (*s*). Local active forces (LA+) and Hi-C–informed interactions (HiC+) are essential to match the experiments. All comparisons here are with IMR90, chr21:28-30Mb data from ref. [8].

### Potential mechanisms underlying extreme chromatin configurations

So far, we have developed a coarsegrained polymer model of chromatin that reproduces 3D distances and their distributions beyond 30 kb. However, the underlying mechanisms at finer scales (nucleosome or kilobase resolution) remain unclear. Since loop extrusion is a well-known active process, one may ask whether extrusion-mediated loop formation (or the lack of it) contribute to these unusual distributions. However, experimental data from [8] in the absence of extrusion, achieved by depleting the cohesin subunit RAD21 using auxin treatment, suggest that loop extrusion does not have a significant impact at the 30 kb scale (Fig. S9). The distributions with and without auxin treatment remain largely unchanged. This is consistent with previous studies showing that, at short length scales (*<* 100 kb), loop extrusion is not the dominant mechanism for loop formation [61].

Hence, going beyond extrusion, we explore other plausible non-equilibrium mechanisms. To understand the nature of the bond-length distribution, we examined whether the probability distribution *P* (*l*_*cg*_) can be expressed as a sum of two or more Gaussian components (see Method S1 E). This is a natural starting point, as Gaussian distributions arise from linear elasticity and thermal fluctuations. We find that *P* (*l*_*cg*_) can be approximately described as a sum of two Gaussians (Fig. 5A). The first Gaussian has a mean near the peak of *P* (*l*_*cg*_) and a width of 72 nm, which is comparable to values obtained from chromatin simulations (Fig. 2A inset). In contrast, the second Gaussian corresponds to a highly extended state, with a mean just below 400 nm and an unusually large width of ≈ 213 nm, exceeding even the standard deviation expected for a random polymer (see Fig. 2A inset). The sum of two Gaussians provides an approximate (though not perfect) fit to the experimental data. For this to occur, two conditions must be satisfied. First, the polymer must transition to a highly extended state, as if intra-chromatin interactions are collectively switched off; such transitions cannot arise from equilibrium thermal fluctuations alone. Second, the unusually large width of this extended state indicates fluctuations stronger than thermal noise, pointing to active, non-equilibrium driving.

**FIG. 5:**
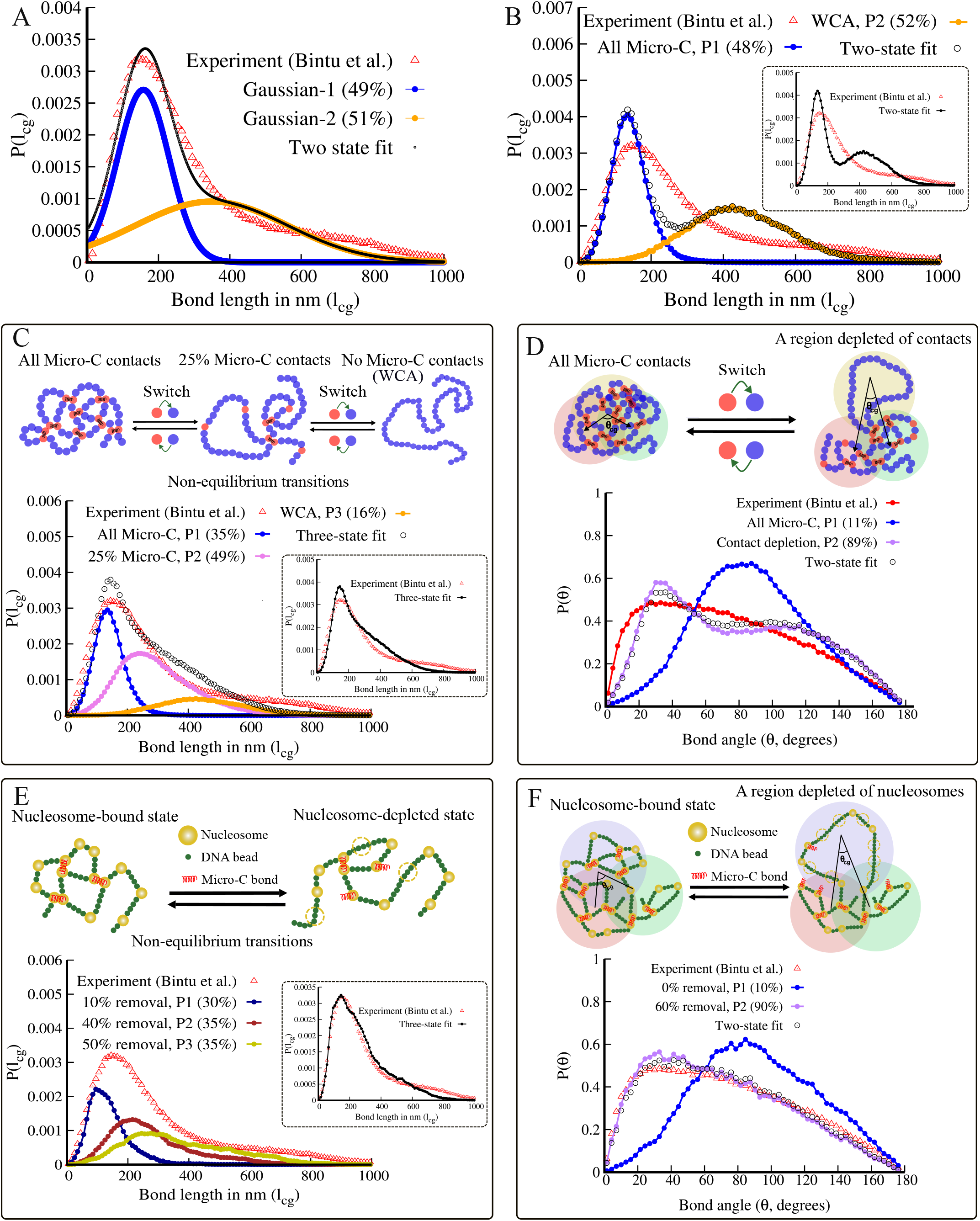
(A) The experimental data can be approximated as a weighted sum of two Gaussians. The weight is given in bracket. The mean (*µ*^*g*^) and standard deviations (*σ*^*g*^) of the Gaussians are 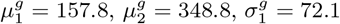, and 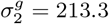 (values rounded). (B) Instead of constructing Gaussians, we take weighted sum of two known polymer distributions. Distributions generated from a folded polymer with all Micro-C interactions as described in the text (P1), and a random self-avoiding polymer (WCA, P2). The best fit (black open circles; also shown in the inset for clarity) is not a perfect match. Top panel: Schematic of chromatin configurations from 3 states: states with all Micro-C contacts(P1), 25% of the Micro-C contacts (P2), and a random (WCA) polymer with no loops (P3). The *l*_cg_ distributions corresponding to these states are in the bottom panel. The best-fitting weighted sum of these distributions is shown in the figure and in inset(black open circles). This is close to the experimental data. (D) To understand angle distributions, we generated configurations by removing Micro-C contacts from 30kb regions keeping the neighboring 30kb intact (top panel). The resulting angle distribution (P2) shifts closer to the experimental data generating acute angles, compared to the null distribution with all micro-C contacts (P1). Here too the best-fitting weighted sum of these distributions is shown in the figure and in inset(black open circles). (E) We simulated chromatin configurations with explicit linker DNA and nucleosomes (top panel). Configurations with different fraction of nucleosome removed generated *l*_cg_ distributions with long tail. The best-fitting weighted sum of these distributions (inset, black open circles) is closer to the experimental data. (F) If nucleosome are removed for a 30kb region, with neighboring 30kb regions intact generates angle distributions similar to what is observed experiments. Here too we computed the best-fitting weighted sum of the distributions. All comparisons here are with IMR90, chr21:28-30Mb data from ref. [8]. To avoid cramped appearance of curves in the figure, we have multiplied *l*_cg_ distributions with the corresponding weights.

To understand this within a polymer framework, we generated two probability distributions from simulated polymer configurations at 200 bp resolution (Fig. 5B). The first distribution (*P*_1_) corresponds to *P* (*l*_*cg*_) obtained from an ensemble of fully folded chromatin configurations that satisfy known inter-nucleosomal contacts from Micro-C maps. The second distribution (*P*_2_) is obtained from configurations of a random self-avoiding (WCA) polymer (SI Text S1 A). We find that a weighted sum of these two distributions (SI Text S1 G) does not adequately reproduce the experimental data (Fig. 5B). We then generated configurations from an intermediate state in which 75% of the Micro-C contacts are removed (Fig. 5C, top panel) and computed the corresponding distribution (SI Text S1 F). A weighted sum of the three distributions provides a much better agreement with the experimental data (Fig. 5C). This suggests that sequential unfolding of chromatin can generate multiple states, and that a mixture of these states can take you close to the observed distribution. This picture is consistent with the coarse-grained model proposed in Fig. 3, where inter-bead distances are extended by non-equilibrium forces. At the molecular level, such unfolding can arise in multiple ways. For example, if nucleosomes collectively switch from an interacting to a non-interacting state (Fig. 5C, top panel), inter-nucleosomal interactions inferred from Micro-C data would be depleted. This may result from non-equilibrium dynamics of histone modifications (see [36]) or fluctuations in DNA-binding proteins. Additionally, condensates of transcription factors, signaling molecules (e.g., ER-*α*, RNA Pol II), and heterochromatin-associated proteins such as HP1 or PRC1/2 are known to compact chromatin locally [28, 63–67]. Transient disassembly of such condensates can lead to local chromatin expansion.

To understand the shift in the angle distribution, we performed the following simulation. We depleted Micro-C contacts from alternate 30 kb regions while keeping contacts in neighboring segments intact (Fig. 5D, top panel). This leads to more acute bending and shifts the angle distribution toward smaller angles (Fig. 5D, bottom panel, SI Text S1 F). A weighted sum of distributions with all contacts and with depleted contacts yields a distribution closer to the experimental data (black open circles) (SI Text S1 G). Although not exact, this result suggests that collective removal of inter-nucleosomal interactions can produce shifts similar to those observed experimentally.

In addition to fluctuations in inter-nucleosome interactions, another possible mechanism is nucleosome disassembly and fluctuations in nucleosome number (Fig. 5E). Remodelers from the SWI/SNF family can disassemble nucleosomes in an ATP-dependent manner. Since nucleosome disassembly releases ∼150 bp of relatively stiff double-stranded DNA, it can lead to extended configurations. We simulated a 90 kb chromatin segment using a high-resolution model with explicit linker DNA and nucleosomes (SI Text S1 H). When a fraction of nucleosomes is removed, the inter-30 kb distance (*l*_*cg*_) distribution develops a long tail. Notably, nucleosome removal leads to distributions with large variance and asymmetry. A weighted combination of states with different fractions of nucleosome removal yields a distribution with a long tail, closer to experimental observations. Furthermore, when a fraction of nucleosomes are removed selectively from the middle 30 kb region while keeping neighboring regions intact, the resulting angle distribution shifts closer to experimental data (Fig. 5F).

Note that we do not claim a unique mechanism for the observed long-tailed distributions; rather, we propose several plausible scenarios that can be tested experimentally, in future. In all cases discussed above, unusually large fluctuations in either inter-nucleosomal interactions or nucleosome density are required. Such collective changes are unlikely to arise from processes obeying detailed balance, as they would require simultaneous disruption of many interactions [36, 62]. This further supports the view that non-equilibrium processes play a key role in generating the observed chromatin configurations.

## Discussion and Conclusion

A quantitative understanding of the polymer properties of human chromatin (stretching and bending elasticities, intra-chromatin interactions) and their coupling to ATP-dependent non-equilibrium processes is essential for simulating and predicting chromatin structure and function. In this context, we discuss the following points to place our results in a broader perspective.

### Polymer properties of human chromatin

In the first section, we presented a “null” model—an equilibrium polymer model with effective interactions derived from Micro-C contact maps. Using our previously developed method [56], we generated an ensemble of nucleosome-resolution configurations consistent with these contacts. This allowed us to estimate key quantities required for coarse-grained human chromatin simulations, including bond-length distributions, spring constants, bending angle distributions, and intra-chromatin interactions (Fig. 1). The predictions of this effective interaction model agree well with the values observed in experiments (Fig. 1H, J, K). However, the model exhibits Gaussian-like elastic behavior, with bond-length distributions centered around a single peak, and fails to capture the large configurational fluctuations and full three-dimensional distance distributions observed experimentally(Fig. 2A-B blue curves, Fig. 3E green curves).

### Chromatin: active polymer with scale-dependent activity

Although there are conceptual similarities between protein folding and chromatin folding, a key distinction is that chromatin operates far from thermal equilibrium; its three-dimensional organization is actively maintained by ATP-consuming molecular machines. However, one of the least-explored aspects of chromatin is the nature of active forces. In this work, we find out the nature and magnitude of the active forces in the scale of 30 kb such that the distance and angle distributions are consistent with experimental observations.

Studies over the past two decades have shown that, at ;S kb scales, various molecular processes generate active forces that drive nucleosome sliding, disassembly, and transcription [60]. Recent studies, as well as our analysis (Fig. S8, Fig. 3), show that loop extrusion, driven by SMC protein complexes, predominantly affects genome organization at length scales *>* 100 kb [61, 68]. In our work here, we predict that active forces also exist at intermediate scales (∼ 10–100 kb), where they manifest as extensile and angular forces that lead to the experimentally observed structural fluctuations. This implies that active forces are inherently scale-dependent, with distinct types of activity operating at different length scales. An open question is whether there are specific molecular processes, analogous to cohesin-mediated loop extrusion, that directly reorganize chromatin at the *∼* 10–100 kb scale. We do not yet know. Our analysis suggests that these intermediate-scale forces may instead emerge from collective effects of molecular processes acting at smaller scales. For example, microphase separation and the assembly and disassembly of condensates are emergent phenomena. Similarly, collective dynamics of histone modifications and DNA-binding proteins can also lead to local chromatin assembly and disassembly.

Our study further shows that directed active forces at one scale can manifest as effective stochastic forces at larger scales. In particular, we find that random active forces at 30 kb scale likely emerge from the averaging of active processes occurring at ∼kb scales. Similarly, the extensile and angular forces identified at intermediate scales become effectively randomized at megabase scales due to averaging over many independent events. Taken together, these considerations suggest that chromatin dynamics is governed by a hierarchy of active forces with distinct characteristics across scales, and that understanding this scale-dependence is essential for a quantitative description of genome organization.

### Why activity is essential to describe the data

Is it essential to invoke an active, non-equilibrium model to describe the phenomena observed here? In principle, one could hypothesize that the long tail in the experimental distance distribution arises from an unconventional effective bond potential between coarse-grained chromatin segments. For example, one could construct a potential of the form *U* ∝ − (*l*_cg_) ln *P* (*l*_cg_) that reproduces the observed distribution. However, such an approach lacks physical consistency within the chromatin polymer framework. Chromatin is composed of interacting nucleosomes, and any realistic attractive interactions among them lead to folded conformations rather than the highly extended configurations observed experimentally.

None of the mechanisms illustrated in Fig. 5 can be realized within an equilibrium framework. The collective opening of chromatin regions necessitates coordinated disassembly of nucleosomes or associated proteins, processes that are inherently non-equilibrium. In addition, the exceptionally large width of the extended-state distribution (Fig. 5A) cannot be explained by thermal fluctuations within any equilibrium polymer model. Taken together, these considerations strongly indicate that non-equilibrium active processes are essential to account for the observed chromatin conformations in a manner that is consistent with the existing knowledge about chromatin.

### Possibility of alternative models

Other than the model we proposed in this work, can there be alternative models to explain the same phenomenon? While it is possible to have alternative models, our work suggests that the key elements of the alternative model has to be non-equilibrium extension and large fluctuations around the extended state. Since we performed Brownian dynamics simulations we adopted a formalism having explicit forces. However, equivalently, one can also use formalism where the chromatin segments would undergo a non-equilibrium switching to an extended state and fluctuate among multiple active states out of equilibrium. Such a model can be simulated using Monte Carlo simulations and fit to the experimental data. Both are equivalent, somewhat like using Langevin equation and Fokker Planck equation to study diffusive motion.

At the molecular level, the inferred active forces likely arise from a combination of processes, including nucleosome remodeling, transcription, histone modification dynamics, and the binding and unbinding of DNA-associated proteins. While our model does not explicitly resolve these microscopic mechanisms, it provides an effective description of their collective impact at the coarse-grained level.

### Chromatin polymer parameters for future simulations

This work provides estimates of polymer parameters and the nature of forces for future simulations of human chromatin. We suggest that the stretching elasticity of CG chromatin segments has two components: an effective harmonic spring contribution and an active stretching contribution. The fluctuation computed from experimental data, quantified as the width of the Gaussian-like peak, is ≈ 96 nm, corresponding to a spring constant of *K*_*s*_ = 0.44 pN/*µ*m (Fig. 2); in dimensionless units, to be used in simulations, this is 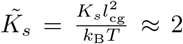. This value is much smaller than those typically used in current polymer simulations. Our results suggest that part of this reduced elasticity arises from active fluctuations. While Fig. 1I provides the spring constant for different coarse-graining scales, our estimate for 30 kb CG segments is *K*_*s*_ = 0.7–2 pN/*µ*m. We recommend using 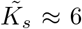 along with active random, extensile and angular forces, as described in this manuscript.

This work also provides new insight into the bending fluctuations of chromatin. Our analysis shows that local chromatin segments explore a wide range of bending angles (0–*π* radians), indicating that chromatin is highly flexible. The distribution deviates significantly from “null” model expectations (Gaussian or *P* ∼ (*θ*) sin *θ* etc) [56] and exhibits a preference for very small intrinsic angles. The broad range of angles suggests that the effective bending stiffness is low, while active angular forces are required to maintain the small intrinsic angle. Analysis of plausible fine-grained models (Fig. 5) suggests an intriguing point: local folding and unfolding of chromatin can modulate the angle distribution, and hence the bendability of coarse-grained chromatin. This indicates that chromatin bending elasticity can be highly non-linear, with local stretching coupled to bending at the coarse-grained scale. A detailed investigation of this coupling is left for future work.

Another important quantity is the intra-chromatin interaction potential. Our work shows that, in coarse-grained models, chromatin segments cannot be treated as hard spheres; instead, they interact via soft repulsive potentials. Fig. 1G provides an estimate of such a potential relevant at sub-megabase scales. This softness arises because long chromatin segments (comprising many nucleosomes) intermix both passively, through diffusion, and actively, through interactions mediated by histone-modifications and proteins as reflected in Micro-C contacts. Although represented as an effective interaction potential, this softness contains an implicit contribution from active processes. This also suggests that at larger scales (*>* 1 Mb), where chromatin segments intermix less, the effective softness is reduced.

### Summary

To conclude, we present a data-driven, non-equilibrium polymer framework that quantitatively captures the structural fluctuations of chromatin polymer. By comparing coarse-grained polymer modeling with existing microscopy data, we show that the large bond-length fluctuations and acute bending angles observed *in vivo* cannot be explained within an equilibrium framework and instead require active, non-equilibrium forces. Our results demonstrate that these active forces are inherently scale-dependent, with directed processes at short length scales giving rise to effective stochastic behavior at larger scales. The proposed model not only reproduces full three-dimensional distance distributions but also enables quantitative inference of effective elastic and active parameters, providing a predictive and experimentally grounded description of chromatin organization. More broadly, this work establishes a general framework for incorporating activity into polymer models and offers a foundation for future studies aimed at linking molecular mechanisms to large-scale genome architecture.

## Methods

### MCCM: MicroC-based Chromatin Model to simulate fine-grained (FG) nucleosome-resolution chromatin

Similar to the method developed in Kadam et al [56] for mESC, chromatin is modeled as a bead–spring polymer, with each bead representing a nucleosome plus linker DNA (200 bp). All neighboring beads along the polymer interact via harmonic potential maintaining the connectivity and any two pair of beads have a WCA (repulsive part of the LJ) potential to prevent overlap (excluded volume interactions). The model then has incorporated intra-chromatin interactions, based on experimentally known Micro-C contacts, as follows. The Micro-C contacts are inserted in two steps. In the first step, only “prominent” contact are inserted. For a given value of genomic separation *s* = |*i* − *j*| all *P*_*ij*_ values are chosen. If contact probability for a pair is one standard deviation larger than the mean for the corresponding *s* value, we define it as prominent contact. That is, *P*_*ij*_ *> P*^*avg*^(*s*) + *σ*^*P*^ (*s*) where *P*^*avg*^(*s*) is the mean contact probability for a given separation s (averaged over all *i* and *j* such that *s* = |*i* −*j*| is a constant), and *σ*^*P*^ (*s*) is the corresponding standard deviation. The selected “prominent pairs” are then connected via harmonic springs to generate one initial configuration of a chromatin polymer given a Micro-C contact map. Several (1000 or above) such initial configurations are generated and all of them are taken to steady-state (“equilibrium” given the interactions) using Langevin polymer simulations (using LAMMPS) generating 1000 independent trajectories. As the second step, for each of these 1000 configurations, non-prominent Micro-C-based contacts are added using a proximity-based interaction method (see SI Text S1 A for details). All these configurations further simulated to obtain an ensemble of steady-state configurations.

### Coarse-graining procedure

Using the MCCM simulations describe above, we generate an ensemble (*>*1000) chromatin configurations at nucleosome resolution. Divide each chromatin configuration of N nucleosomes into segments (sub-polymers) of length *n*_*b*_ nucleosomes. This gives us CG polymer length *N/n*_*b*_ CG beads. Bond length or the distance between two neighboring CG beads (*l*_*cg*_), the bending angle between three consecutive beads (*θ*_*cg*_) and their probability distributions were computed (see Fig. 2A-B). *l*_*cg*_ gave as the effective size (equivalent of diameter) of a CG bead. The inverse of the variance 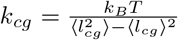 gives us the spring constant of the CG polymer. An effective intra-chromatin interaction between any two non-bonded CG beads of human chromatin was computed using Iterative Boltzmann Inversion (IBI) as described in Kadam et al. [56] (also see S1 B).

### HCCM: HiC-based Chromatin Model to simulate long (∼ Mbp) chromatin as a coarse-grained bead-spring chain

To simulate long region (e.g., chr21:28–30 Mb, IMR90) a CG polymer was constructed with each bead as 5kb (*n*_*b*_ = 25). Every neighboring beads along the polymer interacted with harmonic bonds having spring constant *k*_*cg*_. Any two non-neighboring bead pairs had the following interactions: (a) repulsive part of the soft potential as described in Fig. 1G. (b) Attractive interaction via a harmonic potential based on Hi-C contact probability data at 5kb resolution taken from Rao et al [5] for IMR90 cell line. The method follows the same two-step process described in MCCM: (i) prominent-contact simulation and (ii) proximity-based contact addition. The simulation can be done using Langevin dynamics or Monte Carlo methods. Same procedure is used for any simulation where Hi-C data is the input. Here in this specific case, we used the Metropolis Monte Carlo algorithm, as it is easier to implement intra-chromatin potential, and generated 1000 independent realizations, each run for 10^6^ Monte Carlo steps, with configurations sampled every 10^4^ steps (see SI Text S1 A for details).

### ADCM: Activity-Driven Chromatin Model that incorporates active forces into coarse-grained chromatin polymer simulations

The chromatin polymer was modeled as a coarse-grained chain of *N* beads connected by harmonic springs. The equation of motion for bead *i* is given by Langevin equation

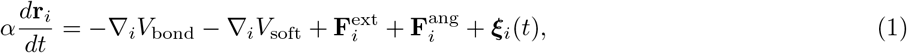

where neighboring beads interact via 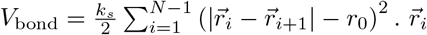 and 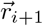 denote the position vectors of the *i*^th^ and *i* + 1^th^ beads respectively, *k*_*s*_ is the spring constant and *r*_0_ is the equilibrium distance between neighboring beads. Each bead represents a coarse-grained chromatin region of 30kb. *V*_soft_ is the repulsive part of the soft-potential as given below (also see [56]),

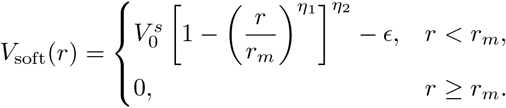

Here, 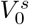 controls the height of the potential at *r* = 0, *r*_*m*_ defines the position of the minimum, and *E* represents the depth of the potential. The parameters *η*_1_ and *η*_2_ can be adjusted to achieve the desired softness. The values of these parameters at 30 kb resolution are derived from Micro-C–based simulation results (see S2) [56, 58].

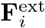 and 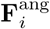 represents extensile and angular active forces along the polymer backbone. To incorporate non-equilibrium activity, we randomly chose three consecutive beads and applied either an extensile force (*F*_ext_) or an angular force (*F*_ang_) as defined below. Then we chose the next three consecutive beads and continued the process such that every bead group had either *F*_ext_ or *F*_ang_ assigned randomly. The microscopy data (Fig. 2) has two features: a single peak and long tail suggesting high extensions with low probability. Since the most probable value (peak) is achievable by the basic model with Micro-C contacts, we hypothesize that the long tail emerge from forces arising from additional non-equilibrium activity. We chose an active force such a way that the probability of having an extensile force *f* is inversely proportional to the strength of the force so that high extension is less probable. Forces were applied to each triplet of consecutive beads (*i* − 1, *i, i*+1), where i is the middle bead of the triplet. The extensile force on each bead is defined as

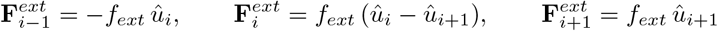

where *f*_*ext*_ is the magnitude of the force and *û*_*i*_ and 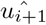 are the unit vectors along the bond as defined below

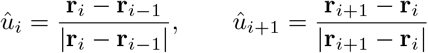

A similar idea was used for the angular force. Microscopy shows that local angles are biased toward smaller values, implying that strong angle-closing (attractive) events should be short-lived. We therefore defined the angular force as

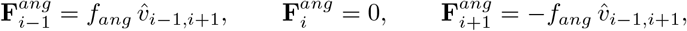

where *f*_*ang*_ is the magnitude of the force, and 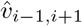 is the unit vector connecting the outer beads of a triplet (i-1 and i+1) as defined below.

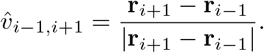

To impliment these forces we did the following procedure: We chose a random number *f*_*r*_ in a range [0.5,5] such that the extensile and the angular forces are given by

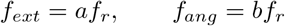

Where, *a* and *b* are constants. To impliment the fact that large forces are less probable, we excerted the force for time a period (Δ*τ*) which is inversely proportional to the strength of the force,

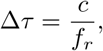

where *c* is a constant. Other than extensile and angular forces, we also apply a random active force *ξ*_*i*_ in the system. It is added as a effective temperature factor in the noise term such that each component obeys 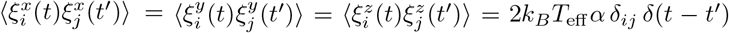 where *α* is a drag coefficient, and *T*_eff_ controls the amplitude of random activity–induced fluctuations [69]. This parameter does not represent a thermodynamic temperature but serves as a measure of nonequilibrium stochastic forcing in the system.

We performed Brownian dynamics simulations of 250 beads polymer where the middle 67 beads represented a 2Mb region of Chr 21 (28-30 Mb) at 30 kb resolution. The integration time step was set to Δ*t* = 0.001, and the spring constant was *k*_*s*_ = 5.9, obtained from fine-grained nucleosome simulation. The simulation length scale was calibrated by matching the peak of the bond-length distribution to the experimentally observed inter-locus separation of 145 nm for 30 kb chromatin segments in IMR90 cells [8]. The optimized parameters used for the active forces were *a* = 28, *b* = 9.5, *T*_eff_ = 15.5 and *c* = 200 (see SI Table S1). Simulations were initialized from random configurations and run for 2 *×* 10^6^ steps using Brownian dynamics implemented in C. For each parameter set, 10 independent trajectories were simulated. System coordinates were recorded every 100 steps, starting after the first 600,000 steps.

### Coarse-grained chromatin polymer simulation with local active forces and HiC-based interactions

The above model has all interactions except attractive interactions between distal segments along the genome. Typically Hi-C contact map represents such interactions often driven by loop extrusion and other intra-chromatin interactions.

We follow similar simulation method as described above and add interaction potentials representing Hi-C contacts. Since Hi-C data at 30 kb resolution were not directly available, we coarse-grained the 10 kb resolution Hi-C matrix from [5]. The raw contact frequency matrix at 10 kb resolution was processed using a 3 *×* 3 sliding window, and the sum of contacts within each window was computed to obtain an effective 30 kb contact matrix. From this coarse-grained matrix, the region of interest (IMR90 chr21: 28–30 Mb) was extracted. To convert contact frequencies into probabilities, we identified the maximum contact value along the diagonal within this region and used it as a normalization factor. All contact values were divided by this factor, resulting in a normalized contact probability matrix. Prominent contacts were defined based on genomic separation *s* = |*i*−*j*|. For each contour distance *s*, we computed the mean contact probability *P*_avg_(*s*) and the corresponding standard deviation *σ*(*s*). A contact between beads *i* and *j* was classified as prominent if 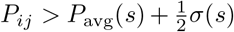. To incorporate structural variability, we generated 1000 independent contact templates. For each prominent contact pair (*i, j*) with probability *P*_*ij*_, a random number *r* ∈ [0, 1] was drawn. The contact was included in a given template if *r < P*_*ij*_. This procedure was repeated for all prominent contacts to generate one template. Repeating this process yielded 1000 distinct contact templates and each template was used to run an independent simulation trajectory. Simulation was performed using overdamped Brownian dynamics. The equation of motion for bead *i* is given by

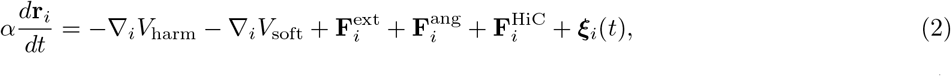

where *V*_harm_ represents the harmonic bond potential, *V*_soft_ denotes non-neighbour bead repulsive interaction and 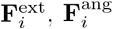, and ***ξ***_*i*_(*t*) are the active forced defined above. Here the maginitude of the soft-potential vary along the contour, as the s increases softness decreases 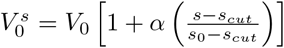. *s*_0_ is the total bead number, and *s*_*cut*_ is the number of CG beads we used in calculating the soft-potential.

We performed Brownian dynamics simulations of a polymer consisting of *N* = 67 beads, using a time step Δ*t* = 0.001. Each trajectory was evolved for a total of 2 *×* 10^7^ steps, and a total of 1000 independent trajectories were generated. After an initial equilibration period, configurations were sampled every 10^4^ steps to construct the ensemble used for analysis.

## Data availability

Relevant data generated from this study are included in this article’s Figures, text, and supplementary information.

## Code availability

All the codes required to perform the simulations are available upon request.

## Acknowledgement

We acknowledge useful discussions with Ajaz Wani, Madan Rao, Srikanth Sastry, PB Sunil Kumar, Gautam Menon, Mandar Inamdar, Vijay Ramani and Shuvadip Dutta. RP acknowledges funding support from ANRF, India.

