## Supplementary text + Figures for "Data-driven polymer modeling reveals how scale-dependent active fluctuations shape chromatin organization"

PACS numbers:

### S1. SUPPLEMENTARY METHODS

#### A. MCCM: Micro-C-based Chromatin Model to simulate a fine-grained (FG) nucleosome-resolution chromatin

We model chromatin as a bead–spring polymer consisting of $N$ beads, where each bead represents a 200 bp segment corresponding to a nucleosome along with linker DNA. Adjacent beads are connected by harmonic springs with stiffness $k_s$. The effective bead diameter is denoted by $\sigma$, which is estimated to be $\sim 20$ nm based on the size of a nucleosome–linker unit (see details below). As input, we use publicly available Micro-C data for human cell lines (HFF and hESC) from Krietenstein et al. [1]. The contact matrix is first KR-normalized and subsequently scaled by the sum of each row to obtain contact probabilities $P_{ij}$.

To generate an ensemble of chromatin configurations consistent with the experimental data, we follow a two-step procedure similar to Kadam et al. [2]. In the first step, we identify a set of prominent contacts from the Micro-C matrix. The prominent contact matrix $P_{ij}^{\text{pr}}$ is defined as

$$P_{ij}^{\text{pr}} = \begin{cases} P_{ij}, & \text{if } P_{ij} > \mu(|i-j|) + \text{s.d.}(|i-j|), \\ 0, & \text{otherwise}, \end{cases} \tag{1}$$

where $\mu(|i-j|)$ and s.d.$(|i-j|)$ denote the mean and standard deviation of contact probabilities for all bead pairs with the same contour separation $|i-j|$. This definition accounts for the inherent dependence of contact probability on genomic separation in polymer systems.

Using $P_{ij}^{\text{pr}}$, we construct an ensemble of 1000 binary contact matrices $C$. For each matrix, the element $C_{ij}$ is assigned as

$$C_{ij} = \begin{cases} 1, & \text{if } r_n < P_{ij}^{\text{pr}}, \\ 0, & \text{otherwise}, \end{cases} \tag{2}$$

where $r_n$ is a uniformly distributed random number in the interval $[0,1]$. Here, $C_{ij} = 1$ indicates the presence of a contact between beads $i$ and $j$.

For each realization of $C$, bead pairs with $C_{ij} = 1$ are connected via harmonic springs with energy

$$E_{\text{mc}} = \frac{k_s}{2} \sum_{i,j>i+1} C_{ij} \left( |\mathbf{r}_i - \mathbf{r}_j| - \sigma \right)^2 . \tag{3}$$

*Electronic address:

Each configuration is then independently equilibrated using Langevin dynamics simulations in LAMMPS [3]. The total energy of the system is given by

$$E = \frac{k_s}{2} \sum_{i=1}^{N-1} \left(|\mathbf{r}_i - \mathbf{r}_{i+1}| - \sigma\right)^2 + \sum_{i,j>i} E_{\mathrm{WCA}}(r_{ij}) + E_{\mathrm{mc}}, \tag{4}$$

where the first term ensures chain connectivity, the second term accounts for excluded volume interactions, and the third term represents Micro-C-derived contacts.

The excluded volume interaction is modeled using the Weeks–Chandler–Andersen (WCA) potential,

$$E_{\mathrm{WCA}}(r) = \begin{cases} 4\epsilon \left[ \left(\frac{\sigma}{r}\right)^{12} - \left(\frac{\sigma}{r}\right)^{6} \right] + \epsilon, & r < 2^{1/6}\sigma, \\ 0, & r \geq 2^{1/6}\sigma. \end{cases} \tag{5}$$

In the second step, we use the equilibrium configurations from the previous step. For each bead pair $i, j$, spring contacts ($C_{ij} = 1$) were assigned to a fraction $P_{ij}$ of configurations with the smallest 3D separations $r_{ij}$. The system was then equilibrated using the updated contact matrix $C$, following the same protocol as in the first step.

We performed Langevin dynamics simulations using LAMMPS for a polymer of size $N = 1000$ (corresponding to a 200 kb region) with spring constant $k_s = 10$ [2]. The initial polymer configuration is generated as a self-avoiding walk. To avoid large forces arising from long-range contacts, we introduce Micro-C-derived interactions progressively. We first include local contacts with $|i - j| < 5$ and equilibrate the system for $10^5$ steps. Subsequently, contacts with increasing contour separations ($|i-j| < 10$, $|i-j| < 25$, etc.) are added in stages, with equilibration performed after each addition. Once all contacts are incorporated, the system is equilibrated for $6 \times 10^6$ time steps. Configurations are sampled every 50,000 steps during the final $3 \times 10^6$ steps for analysis.

**HCCM: Hi-C-based Chromatin Model for simulating long ($\sim$ Mbp) chromatin as a coarse-grained bead–spring chain:** In this model, chromatin is represented as a polymer comprising $N$ beads, where each bead corresponds to a 5 kb segment of chromatin. The bond-length distribution and corresponding spring constants are obtained from Fig. 1 H-I. Non-neighbouring beads interact via a soft potential (see Fig. 1G), derived using the iterative Boltzmann inversion (IBI) procedure (see below). The overall simulation protocol is identical to that used in MCCM and follows a two-step procedure [2]. As input, we use publicly available Hi-C data for the IMR90 cell line from Rao et al. [4]. We performed Monte Carlo (MC) simulations using the Gillespie algorithm implemented in C. We simulate a 400-bead polymer corresponding to a 2 Mb genomic region (IMR90, chr21: 28–30 Mb). Each simulation is run for a total of $10^6$ steps. After an initial equilibration phase, configurations are sampled every $10^4$ steps for subsequent analysis.

**SAW polymer (no Micro-C contacts):** As a control, we performed the same simulations for a self-avoiding walk (SAW) polymer. We simulated $N = 1000$ beads, where each bead represents 200 bp of chromatin. The simulation method is the same as in the first part of MCCM (see above), without Micro-C-derived contacts, by setting $C_{ij} = 0$ for all bead pairs. The total energy is given by

$$E = \sum_{i=1}^{N-1} \frac{k_s}{2} \left(|\mathbf{r}_i - \mathbf{r}_{i+1}| - \sigma\right)^2 + \sum_{i=1}^{N} \sum_{j>i} E_{\mathrm{WCA}}\left(|\mathbf{r}_{ij}|\right). \tag{6}$$

Here, $E_{\mathrm{WCA}}$ accounts for the excluded volume interactions and is modeled using the Weeks–Chandler–Andersen (WCA) potential (see above). The simulated equilibrium configurations are coarse-grained (see the coarse-graining procedure section in the main text) to 30 kb, and the bond-length distribution ($P_{\mathrm{lcg}}$) is calculated (see Fig. 5 A). The average bond length at 30 kb resolution is plotted as a dashed line in Fig. 2A.

**Ideal Gaussian chain:** The ideal Gaussian chain comprises $N$ beads connected by harmonic springs in the absence of excluded volume interactions ($E_{\mathrm{WCA}} = 0$), with spring constant $k_s = 10$. The total energy of the system is given by

$$E = \sum_{i=1}^{N-1} \frac{k_s}{2} \left(|\mathbf{r}_i - \mathbf{r}_{i+1}| - \sigma\right)^2. \tag{7}$$

This model serves as a reference for comparing the width (see below) of bond-length distributions obtained from microscopy and simulations.

#### B. Iterative Boltzmann Inversion Method

To determine the effective nonbonded interaction potentials at different coarse-graining levels, we used the nucleosome-resolution simulation data for the HFF cell line corresponding to the genomic region chr21:28–28.2 Mb. From the simulated configurations, we computed the distribution of distances between nonbonded beads for different coarse-grained resolutions (2 kb, 5 kb, 10 kb, 30 kb, and 50 kb). These distributions serve as the target distributions, denoted as $P_{\text{target}}(r)$, for the corresponding coarse-grained models.

For a given coarse-graining level, the polymer consists of $N/n_b$ beads, where $n_b$ is the number of fine-grained units per coarse-grained bead. The bonded interactions between consecutive beads are modeled using a harmonic spring potential with spring constant $K_{\text{cg}}$ (see Fig. 1I) and equilibrium bond length $l_{\text{cg}}$ (see Fig. 1H). We run a Monte Carlo simulation, and the total energy of the system is given by

$$E = \sum_i \frac{K_{\text{cg}}}{2} \left(|\mathbf{r}_i - \mathbf{r}_{i+1}| - l_{\text{cg}}\right)^2 + \sum_{i,j>i+1} V^{\text{nb}}(|\mathbf{r}_i - \mathbf{r}_j|), \tag{8}$$

where $V^{\text{nb}}(r)$ represents the nonbonded interaction potential, which is obtained using the iterative Boltzmann inversion (IBI) scheme [2]. We initialize the potential as $V_0^{\text{nb}}(r) = 0$ and perform Metropolis Monte Carlo simulations to sample equilibrium configurations. After equilibration, the distance distribution between nonbonded beads, $P_i(r)$, is computed at iteration $i$. The convergence of the method is monitored using the Kullback–Leibler (KL) divergence between the target and simulated distributions:

$$D_{\text{KL}}(P_{\text{target}} \,\|\, P_i) = \sum_r P_{\text{target}}(r) \ln\left(\frac{P_{\text{target}}(r)}{P_i(r)}\right). \tag{9}$$

If convergence is not achieved, the nonbonded potential is updated according to

$$V_{i+1}^{\text{nb}}(r) = V_i^{\text{nb}}(r) + \alpha(r) k_B T \ln\left(\frac{P_i(r)}{P_{\text{target}}(r)}\right), \tag{10}$$

where $k_B T$ is the thermal energy. The function $\alpha(r)$ is chosen as a decaying function,

$$\alpha(r) = 0.2\, e^{-r^2/2}, \tag{11}$$

to ensure that the resulting potential remains short-ranged. This iterative procedure is repeated until the KL divergence approaches zero, indicating convergence of the simulated distribution to the target distribution.

The final converged nonbonded potential obtained from the IBI procedure is numerical in nature. To obtain a smooth and transferable representation, we fit this potential to an analytic functional form given by

$$V_{\text{soft}}(r) = \begin{cases} V_0 \left[1 - \left(\dfrac{r}{r_m}\right)^{\eta_1}\right]^{\eta_2} - \epsilon, & r < r_m, \\ \dfrac{1}{2}\epsilon \left[\cos(\mu r^2 + \nu) - 1\right], & r_m \le r < r_c, \\ 0, & r \ge r_c. \end{cases} \tag{12}$$

The short-range regime ($r < r_m$) corresponds to the repulsive component of the interaction [5], whereas the intermediate regime ($r_m \le r < r_c$) describes the attractive part of the potential [6]. The parameters $\mu$ and $\nu$ are obtained by imposing continuity and smoothness constraints, given by

$$\mu r_m^2 + \nu = \pi, \quad \mu r_c^2 + \nu = 2\pi. \tag{13}$$

These conditions ensure that the potential attains a value of $-\epsilon$ at $r = r_m$ and continuously approaches zero at the cutoff distance $r = r_c$. The corresponding fitted parameters for different coarse-graining levels are summarized in Table S2.

#### C. Size of a 200 bp chromatin bead

The size of a 200 bp chromatin bead ($\sigma$) is expected to exceed that of a single nucleosome ($\sim$ 11 nm). Adjacent nucleosomes, connected by a $\sim$ 50 bp linker DNA, are typically separated by distances up to $\sim$ 28 nm [2]. In contrast, nucleosomes that are not directly connected can approach each other more closely, with separations of $\sim$ 11–12 nm due to histone tail interactions and associated protein binding. Taking these limits into account, an effective bead size of $\sim$ 20 nm is a reasonable estimate. This value also reflects the in vivo scenario, where nucleosomes are often associated with various proteins (e.g., acetyltransferases, methyltransferases, HMG proteins, HP1, and chromatin remodelers), effectively increasing their occupied volume. Moreover, this estimate is consistent with previous studies reporting $\sigma \approx 25$ nm for 250 bp chromatin beads [7].

#### D. Optimization of Activity Parameters

Activity parameters were obtained by matching the bond-length and bond-angle distributions ($P(l_{\rm cg})$ and $P(\theta_{\rm cg})$ ) from simulations to those from experiment. Target probability distributions for bond length and bond angle, $P_{\rm exp}(r)$ and $P_{\rm exp}(\theta)$, were calculated from Bintu et al. super-resolution microscopy data at 30 kb resolution [8]. For a given parameter set $(a, b, T_{\rm eff})$, active polymer dynamics were simulated to generate ensembles of chromatin configurations. From the resulting trajectories, the corresponding simulated distributions $P_{\rm sim}(r; a, b, T_{\rm eff})$ and $P_{\rm sim}(\theta; a, b, T_{\rm eff})$ were constructed using identical binning and normalization as the experimental data. The agreement between simulation and experiment was quantified using the Jensen–Shannon divergence (JSD), computed separately for bond-length and bond-angle distributions and summed to define the total discrepancy. The JSD between two probability distributions $P$ and $Q$ is defined as

$$JSD(P \,\|\, Q) = \frac{1}{2}\sum_i P(i)\log\frac{P(i)}{M(i)} + \frac{1}{2}\sum_i Q(i)\log\frac{Q(i)}{M(i)}, \quad M(i) = \frac{1}{2}\big(P(i) + Q(i)\big). \tag{14}$$

The total divergence is then given by

$$JSD_{\rm total}(a, b, T_{\rm eff}) = JSD(P_{\rm sim}(r), P_{\rm exp}(r)) + JSD(P_{\rm sim}(\theta), P_{\rm exp}(\theta))\,. \tag{15}$$

A structured grid search was performed over the activity parameter space, and for each parameter set the total divergence $JSD_{\rm total}$ was evaluated. The optimal activity parameters were identified by solving $(a^*, b^*, T^*_{\rm eff}) = \arg\min_{a,b,T_{\rm eff}}\; JSD_{\rm total}(a, b, T_{\rm eff})$. The optimal activity parameters with corresponding JSD values for different cell lines are shown in Table S1.

#### E. Two-Gaussian fitting of the bond-length distribution:

To obtain the best two-Gaussian fit to the experimental $l_{\rm cg}$ distribution (IMR90, chr21: 28–30 Mb), we model the probability distribution ($P_{\rm fit}(x)$) as a weighted sum of two Gaussian components:

$$P_{\rm fit}(x) = w\, g_1(x) + (1 - w)\, g_2(x), \tag{16}$$

where $w \in [0, 1]$ is the mixing weight, and $g_1(x)$ and $g_2(x)$ are normalized Gaussian distributions given by

$$g_i(x) = \frac{1}{\sqrt{2\pi}\,\sigma_i}\exp\left(-\frac{(x - \mu_i)^2}{2\sigma_i^2}\right), \quad i = 1, 2. \tag{17}$$

We performed a grid search over the parameter space $(\mu_1, \mu_2, \sigma_1, \sigma_2, w)$. The means $\mu_1$ and $\mu_2$ were varied uniformly over the range of the data, $\mu \in [x_{\rm min}, x_{\rm max}]$, using 40 equally spaced points. The standard deviations $\sigma_1$ and $\sigma_2$ were varied over the range $\sigma \in \left[\frac{x_{\rm max}-x_{\rm min}}{50}, \frac{x_{\rm max}-x_{\rm min}}{5}\right]$, sampled using 20 equally spaced points. Here, $x_{\rm min}$ and $x_{\rm max}$ denote the minimum and maximum values of $x$ in the experimental distribution $P_{\rm exp}(x)$. The mixing weight was varied over $w \in [0, 1]$ using 50 equally spaced values. For each parameter combination, the model distribution $P_{\rm fit}(x)$ was evaluated at the discrete data points, and the deviation from the experimental distribution ($P_{\rm exp}(x)$) was quantified using the sum of squared errors:

$$\text{Error} = \sum_i \left[P_{\rm exp}(x_i) - P_{\rm fit}(x_i)\right]^2. \tag{18}$$

The optimal parameters $(\mu_1, \mu_2, \sigma_1, \sigma_2, w)$ were identified as those that minimize this error. Using these best-fit parameters, the individual Gaussian components and the total fitted distribution were reconstructed for comparison with the experimental data (see Fig. 5A).

#### F. Simulating chromatin polymer with reduced Micro-C contacts

**Simulating chromatin polymer with randomly reduced Micro-C contacts:** In Fig. 5C, we simulated a chromatin polymer using prominent Micro-C contacts from the HFF cell line (chr21: 28–28.2 Mb) at 200 bp resolution, based on experimental data from Krietenstein et al. [1]. The simulation procedure was identical to that described in the first part of MCCM (see above). From the set of prominent contacts, we randomly removed $p$% of contacts, where $p$ varied from 0 to 75, and performed simulations using the reduced contact sets (see MCCM method for details). In Fig. 5C, we show three cases: all Micro-C contacts (0% removal), 25% of the Micro-C contacts retained (i.e., 75% removal), and a self-avoiding walk (SAW) polymer (described above). The resulting structures were coarse-grained to a 30 kb resolution, and the bond-length distribution $P(l_{\rm cg})$ was computed.
**Simulating chromatin polymer with regionally reduced Micro-C contacts:** In Fig. 5D, we simulated a chromatin polymer using the same genomic region and resolution as described above. The simulation procedure was identical to that described in the first part of MCCM (see above). The 200 kb region was divided into 30 kb segments, and from every alternate segment, all prominent contacts within that segment, as well as any contacts involving that region, were removed, resulting in alternating 30 kb contact-depleted regions. We then simulated 1000 independent trajectories as described in the MCCM method (see above). From the equilibrium configurations, the system was coarse-grained to a 30 kb resolution, and $P(\theta_{\rm cg})$ was computed.

#### G. Linear combination fitting of experimental distributions using simulation-derived distributions:

In Fig. 5, we fit the experimental bond-length and angle distributions ($P(l_{\rm cg})$ and $P(\theta_{\rm cg})$ ), obtained from Bintu et al. [8], using distributions derived from our simulations. Specifically, the experimental probability distribution $P_{\rm exp}(x)$ ($P(l_{\rm cg})$ or $P(\theta_{\rm cg})$ ) is modeled as a weighted linear combination of simulation-derived distributions.

For a two-state model (Fig. 5D and F), the experimental distribution is expressed as

$$P_{\rm model}(x) = w_1 P_1(x) + w_2 P_2(x), \tag{19}$$

subject to

$$w_1 + w_2 = 1, \quad w_1, w_2 \in [0, 1]. \tag{20}$$

The weight $w_1$ is varied uniformly over the interval $[0, 1]$ using 100 equally spaced points, and $w_2$ is set as $w_2 = 1 - w_1$.

For a three-state model (Fig. 5C and E), we write

$$P_{\rm model}(x) = w_1 P_1(x) + w_2 P_2(x) + w_3 P_3(x), \tag{21}$$

with

$$w_1 + w_2 + w_3 = 1, \quad w_i \in [0, 1]. \tag{22}$$

The weight $w_1$ and $w_2$ is varied uniformly over the interval $[0, 1]$ using 50 equally spaced points, and $w_3$ is set as $w_3 = 1 - w_1 - w_2$.

For each set of weights, the model distribution is evaluated at the discrete data points $\{x_i\}$, and the deviation from the experimental distribution is quantified using the sum of squared errors:

$$\text{Error} = \sum_i \left[P_{\rm exp}(x_i) - P_{\rm model}(x_i)\right]^2 . \tag{23}$$

The optimal weights are chosen as those that minimize this error.

#### H. Nucleosome–Linker Model with Prominent Micro-C Contacts

Chromatin is modeled as a bead–spring polymer consisting of two types of spherical beads: linker DNA (D) and nucleosomes (Nu). Each DNA bead represents 10.5 base pairs (bp) and has a diameter $\sigma$ ($\sim$ 3.57nm), which defines the fundamental unit of length in the system. Each nucleosome bead represents a histone octamer wrapped with 147 bp of DNA and is assigned a diameter of $3\sigma$. At least one DNA bead is placed between consecutive nucleosomes.

Adjacent beads are connected by harmonic springs with stiffness $k_s = 100\, k_\mathrm{B}T/\sigma^2$. The harmonic potential, $\mathrm{V_{bond}(r)}$ is defined as

$$V_\mathrm{bond}(r) = \frac{k_s}{2}(r - r_0)^2, \tag{24}$$

where $r$ is the bond length and $r_0$ is the equilibrium bond distance. The equilibrium distances are set to $\sigma$ for DNA–DNA bonds, $2\sigma$ for DNA–Nu bonds, and $6\sigma$ for Micro-C bonds. Chain stiffness is incorporated using a cosine bending potential:

$$V_\mathrm{bend}(\theta) = k_b\left[1 - \cos(\theta - \theta_0)\right], \tag{25}$$

where $\theta$ is the bond angle. For DNA–DNA–DNA triplets, the parameters are $(\theta_0, k_b) = (180°, 28\, k_\mathrm{B}T)$, while for DNA–Nu–DNA triplets, $(\theta_0, k_b) = (60°, 3.8\, k_\mathrm{B}T)$ [9]. The total potential energy of the system is given by:

$$E = \sum_{i=1}^{N-1} \frac{k_s}{2}\left(|\mathbf{r}_i - \mathbf{r}_{i+1}| - r_0\right)^2 + \sum_{i=1}^{N-2} k_b\left[1 - \cos(\theta_i - \theta_0)\right] + \sum_{i<j} E_\mathrm{WCA}(|\mathbf{r}_i - \mathbf{r}_j|) + E_\mathrm{mc}. \tag{26}$$

where $\mathrm{E_{WCA}}$ represents excluded volume interactions and is modeled using the Weeks–Chandler–Andersen (WCA) potential with $\epsilon = k_\mathrm{B}T$ and a cutoff distance of $2^{1/6}\sigma$. $E_\mathrm{mc}$ represents Micro-C derived contact as mentioned before [1]. Only prominent contacts $P^r_{ij}$ are selected for interaction (similar to the first part of MCCM). The polymer was initialized as a self-avoiding walk. Micro-C bonds were then introduced progressively, starting from short genomic separations and extending to longer ones, with each stage followed by equilibration over $10^4$ time steps. After incorporating all contacts, the system is further equilibrated for $2 \times 10^7$ steps. Configurations are sampled every $10^4$ steps during the final $10^7$ steps, yielding a total of $10^3$ configurations. We simulated a 90 kb chromatin region (HFF cell line, chr17:50.165-50.255Mb) using Langevin dynamics in LAMMPS, with shrink-wrapped boundaries and in reduced Lennard–Jones units.

**Simulating nucleosome-depleted state:** To generate a nucleosome-depleted state (see Fig. 5E), we randomly removed $p\%$ of nucleosomes from the 90 kb chromatin polymer, where $p$ ranges from 10 to 50. Upon nucleosome removal, the associated 147 bp of wrapped DNA was assumed to unwrap and modeled as linker DNA, represented by 14 DNA beads represented by 14 DNA beads (10.5 bp per bead). The resulting configurations were then equilibrated using the simulation protocol described above. We performed 100 independent trajectories, each initialized from an independent starting configuration. The system was subsequently coarse-grained to a 30 kb resolution, and the bond-length distribution ($P(l_\mathrm{cg})$ ) was computed.

**Simulating a region depleted of nucleosomes:** As described above, a fraction of nucleosomes was removed; however, in this case, the removal was restricted to the central 30 kb of the 90 kb chromatin polymer rather than applied uniformly. Specifically, $p\%$ of nucleosomes were removed from the middle 30 kb region, where $p$ ranges from 0 to 60. The resulting configurations were equilibrated using the same simulation method. From the equilibrium configurations, the system was coarse-grained to a 30 kb resolution, and the bond-angle distribution was computed (see Fig. 5F).

### I. Quantities measured

• **Contact probability:** The contact probability between a pair of beads $i$ and $j$ is calculated as

$$P_{ij} = \frac{1}{N_\mathrm{conf}} \sum_{k=1}^{N_\mathrm{conf}} H(r_\mathrm{cut} - r_{ij}^{(k)}), \tag{27}$$

where $N_\mathrm{conf}$ is the total number of sampled polymer configurations, $H(x)$ denotes the Heaviside step function, and $r_{ij}^{(k)}$ is the distance between beads $i$ and $j$ in the $k$-th configuration. Two beads are considered to be in contact if their separation is less than a cutoff distance $r_\mathrm{cut}$. For the nucleosome-level fine-grained model, the contact cutoff is set to $r_\mathrm{cut} = 1.2\sigma$ (Fig. 1B). For the coarse-grained simulations, we employ $r_\mathrm{cut} = 1.0\sigma$ for Fig. 1J and $r_\mathrm{cut} = 1.3\sigma$ for Fig. 4C. Note that these simulations are at different resolutions. Simulations in Fig. 1J and Fig. 4C are performed with a soft potential, and the cutoff distance for the soft potential is an unknown parameter. However, the main prediction of the simulation is the 3D distance, which is independent of the cutoff parameter.

• **Bond length.** The coarse-grained bond length, $l_\mathrm{cg}$, is defined as the distance between consecutive coarse-grained (CG) units. Its computation depends on whether the coordinates arise from experiments or simulations.

**(i) Experimental data (microscopy).** For microscopy data, 3D coordinates of 30 kb genomic segments are directly available. The bond length is computed as

$$l_i^{\mathrm{cg}} = |\mathbf{r}_i - \mathbf{r}_{i+1}|\,, \tag{28}$$

where $\mathbf{r}_i$ and $\mathbf{r}_{i+1}$ denote the positions of consecutive genomic segments.

**(ii) Simulation data.** For simulation data, two scenarios arise depending on the resolution.

*(a) Coarse-grained simulations.* If the simulation is already at the coarse-grained level (one bead represents a 30 kb segment), the bond length is computed directly as

$$l_i^{\mathrm{cg}} = |\mathbf{r}_i - \mathbf{r}_{i+1}|\,. \tag{29}$$

*(b) Fine-grained simulations.* If the simulation is at a finer resolution, CG beads are constructed by grouping underlying units ($\mathrm{n_b}$ fine grained beads), and their positions are defined via the center of mass. The bond length is then given by

$$l_i^{\mathrm{cg}} = \left|\mathbf{r}_i^{\mathrm{com}} - \mathbf{r}_{i+1}^{\mathrm{com}}\right|, \tag{30}$$

where $\mathbf{r}_i^{\mathrm{com}}$ denotes the center-of-mass position of the $i$-th CG bead.

• **Bond angle.** The coarse-grained bond angle, $\theta_i^{\mathrm{cg}}$, is defined as the internal angle formed by three consecutive coarse-grained (CG) units. The angle is computed from the vectors connecting adjacent beads using the dot product.

**(i) Experimental data (microscopy).** For microscopy data, the 3D coordinates of 30 kb genomic segments are directly available. The bond angle is computed as

$$\theta_i^{\mathrm{cg}} = \cos^{-1}\left(\frac{(\mathbf{r}_{i-1} - \mathbf{r}_i)\cdot(\mathbf{r}_{i+1} - \mathbf{r}_i)}{|\mathbf{r}_{i-1} - \mathbf{r}_i|\,|\mathbf{r}_{i+1} - \mathbf{r}_i|}\right), \tag{31}$$

where $\mathbf{r}_{i-1}$, $\mathbf{r}_i$, and $\mathbf{r}_{i+1}$ denote the positions of three consecutive genomic segments.

**(ii) Simulation data.** For simulation data, the definition depends on the resolution.

*(a) Coarse-grained simulations.* If the simulation is already at the coarse-grained level, the bond angle is computed directly from bead positions as

$$\theta_i^{\mathrm{cg}} = \cos^{-1}\left(\frac{(\mathbf{r}_{i-1} - \mathbf{r}_i)\cdot(\mathbf{r}_{i+1} - \mathbf{r}_i)}{|\mathbf{r}_{i-1} - \mathbf{r}_i|\,|\mathbf{r}_{i+1} - \mathbf{r}_i|}\right). \tag{32}$$

where $\mathbf{r}_i$ represents the position vector of the $i^{th}$ CG bead.

*(b) Fine-grained simulations.* If the simulation is at a finer resolution, CG bead positions are obtained via center-of-mass mapping of the $\mathrm{n_b}$ fine grained beads. The bond angle is then computed as

$$\theta_i^{\mathrm{cg}} = \cos^{-1}\left(\frac{(\mathbf{r}_{i-1}^{\mathrm{com}} - \mathbf{r}_i^{\mathrm{com}})\cdot(\mathbf{r}_{i+1}^{\mathrm{com}} - \mathbf{r}_i^{\mathrm{com}})}{|\mathbf{r}_{i-1}^{\mathrm{com}} - \mathbf{r}_i^{\mathrm{com}}|\,|\mathbf{r}_{i+1}^{\mathrm{com}} - \mathbf{r}_i^{\mathrm{com}}|}\right), \tag{33}$$

where $\mathbf{r}_i^{\mathrm{com}}$ denotes the center-of-mass position of the $i$-th CG bead.

• **Spring constant.** We model the interaction between consecutive coarse-grained (CG) beads using harmonic springs. The effective spring constant for a given coarse-graining level ($n_b$) is computed as

$$K_{\mathrm{cg}} = \frac{k_B T}{\langle l_{cg}^2\rangle - \langle l_{cg}\rangle^2}. \tag{34}$$

where $k_B T$ is the thermal energy, and $l_{cg}$ denotes the CG bond-length.

• **Stratum-adjusted correlation coefficient (SCC).** To quantify the similarity between experimental and simulated contact maps, we compute the stratum-adjusted correlation coefficient (SCC) (Yang et al. [10]). Prior to SCC calculation, both contact maps are smoothed using a two-dimensional mean filter, where each element is replaced by the average over a square neighborhood of size $(2h+1)\times(2h+1)$ (a square of size 2h + 1 centered around $P_{ij}$ in the contact matrix). The data are then stratified based on genomic separation $|i-j|$, and for each stratum $k$, we compute the Pearson correlation coefficient between the logarithm of contact probabilities, $\log P_{ij}$, obtained from experiment and simulation. The final SCC is calculated as a weighted average of the stratum-wise Pearson correlations. In this work, we use $h = 5$ for smoothing; the number of strata is set to $k = 200$ for Fig. 1B and J, and $k = 20$ for Fig. 4C.

• **Pearson correlation.** To quantify the agreement between simulated and experimentally measured chromatin organization, we compute the Pearson correlation coefficient between the pairwise 3D distances obtained from simulations and microscopy. Specifically, for each pair of genomic segments, the predicted distance from simulations was compared to the corresponding microscopy distance in a one-to-one manner. The Pearson correlation coefficient, $r_p$, is defined as

$$r_p = \frac{\sum_i (d_i^{\rm sim} - \bar{d}^{\rm sim})(d_i^{\rm exp} - \bar{d}^{\rm exp})}{\sqrt{\sum_i (d_i^{\rm sim} - \bar{d}^{\rm sim})^2}\sqrt{\sum_i (d_i^{\rm exp} - \bar{d}^{\rm exp})^2}}, \quad (35)$$

where $d_i^{\rm sim}$ and $d_i^{\rm exp}$ denote the simulated and experimental distances for the $i$-th pair, respectively, and $\bar{d}^{\rm sim}$ and $\bar{d}^{\rm exp}$ are their corresponding means. The correlation was computed using standard Python libraries.

• **Width estimation from bond-length distribution.** To quantify the width of the bond-length distribution, we compute a generalized full-width measure based on a fixed fraction of the peak height (Fig. 2A). From the normalized probability density $P(r)$, the peak value $P_{\rm max}$ and its corresponding position $r_0$ are first identified. We define the width at approximately 88% of the peak height, i.e., at $P(r) = P_{\rm max}\exp(-1/8)$ The left ($r_-$) and right ($r_+$) crossing points are determined by linearly interpolating between adjacent histogram bins around this threshold. The resulting width is then computed as $w = r_+ - r_-$. This choice corresponds to a Gaussian-equivalent measure of width and is less sensitive to noise in the distribution tails.

### S2. SUPPLEMENTARY TABLES

| Cell line | $T_{\rm eff}$ | $a$ (JSD in $10^{-3}$) | $b$ (JSD in $10^{-3}$) | JSD ($10^{-3}$) |
|---|---|---|---|---|
| IMR90 | 15.5 | 28.0 (3.06) | 9.5 (0.736) | 3.804 |
| A549 | 12.0 | 29.0 (2.05) | 5.0 (0.178) | 2.233 |
| HCT116 | 11.0 | 29.0 (2.90) | 6.0 (0.401) | 3.306 |

**TABLE S1:** Activity parameters obtained for different cell lines for the chromatin region chr21:28–30 Mb at 30kb resolution, along with the corresponding JSD values.

| CG Size | Soft Potential Parameters | | | | | | | |
|---|---|---|---|---|---|---|---|---|
| | $V_0$ | $\epsilon$ | $r_m$ | $\eta_1$ | $\eta_2$ | $r_c$ | $\mu$ | $\nu$ |
| 2kb (n=10) | 3.59 | 0.17 | 1.35 | 1.94 | 4.86 | 2.14 | 1.05 | 1.44 |
| 5kb (n=25) | 2.79 | 0.24 | 1.35 | 1.91 | 4.00 | 2.31 | 0.71 | 2.48 |
| 10kb (n=50) | 2.27 | 0.36 | 1.35 | 2.51 | 4.61 | 2.33 | 0.80 | 1.91 |
| 30kb (n=150) | 2.75 | 0.99 | 1.35 | 2.26 | 3.03 | 2.26 | 0.91 | 1.80 |
| 50kb (n=250) | 2.86 | 1.93 | 1.35 | 2.37 | 3.62 | 2.40 | 0.75 | 2.20 |

**TABLE S2:** Derived coarse-grained (CG) soft-potential polymer parameters from Micro-C simulations for the HFF cell line, corresponding to the chr21:28–28.2 Mb genomic region.

### S3. SUPPLEMENTARY FIGURES

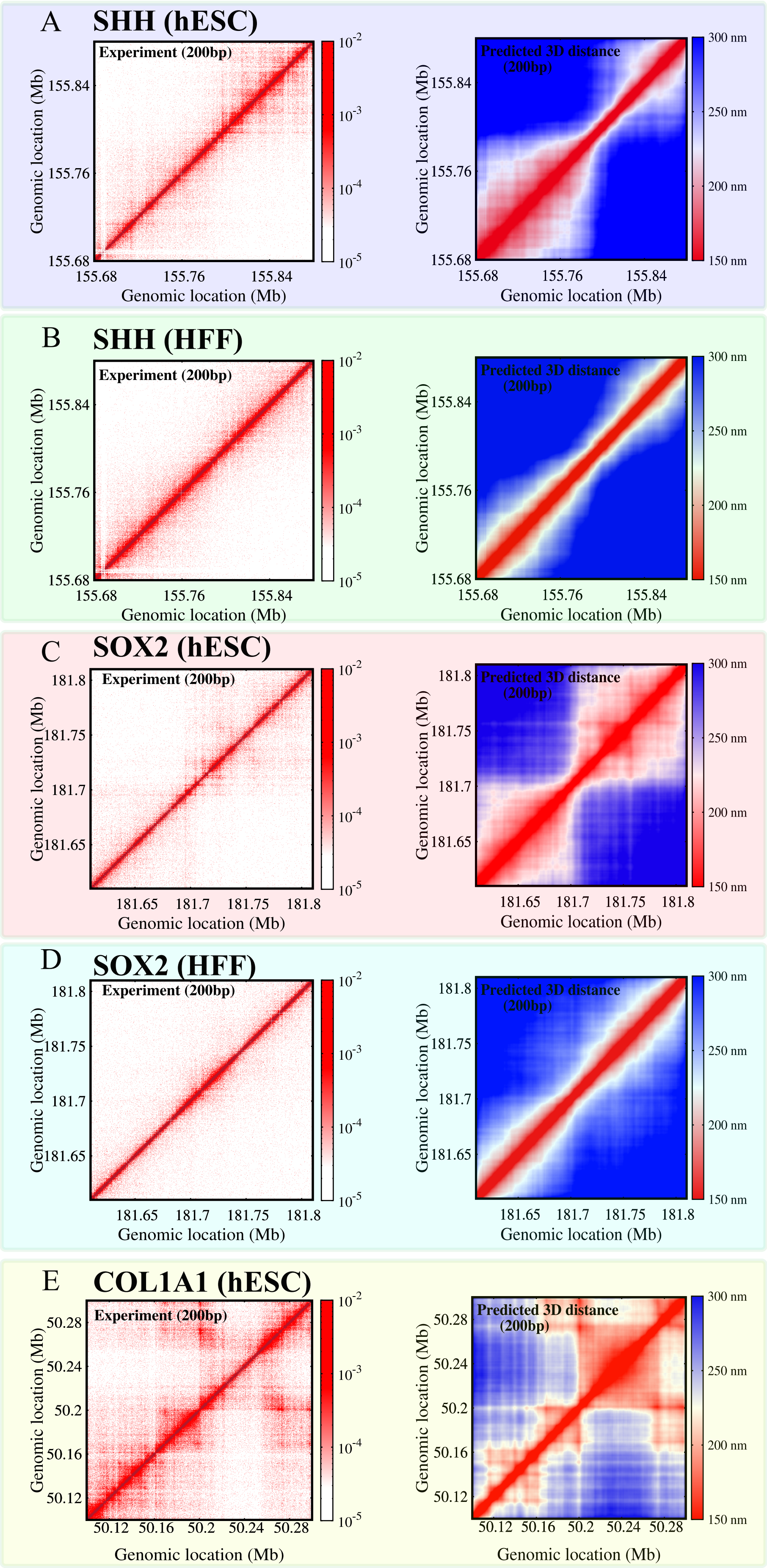


**FIG. S1:** Comparison between Micro-C contact maps (left) and the corresponding predicted 3D distance maps from the MCCM simulation (right) for three genomic regions: SHH (A, B), SOX2 (C, D), and COL1A1 (E).

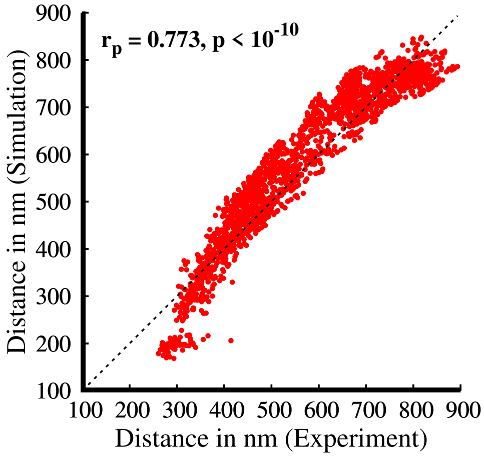


**FIG. S2:** One-to-one comparison of predicted 3D distances from Fig. 1. The x-axis represents the average microscopy-measured distance, while the y-axis shows the corresponding average distance obtained from simulations. Data are shown for the IMR90 cell line at Chr21:28–30 Mb. Simulated configurations at 5 kb resolution were coarse-grained to 30 kb for direct comparison.

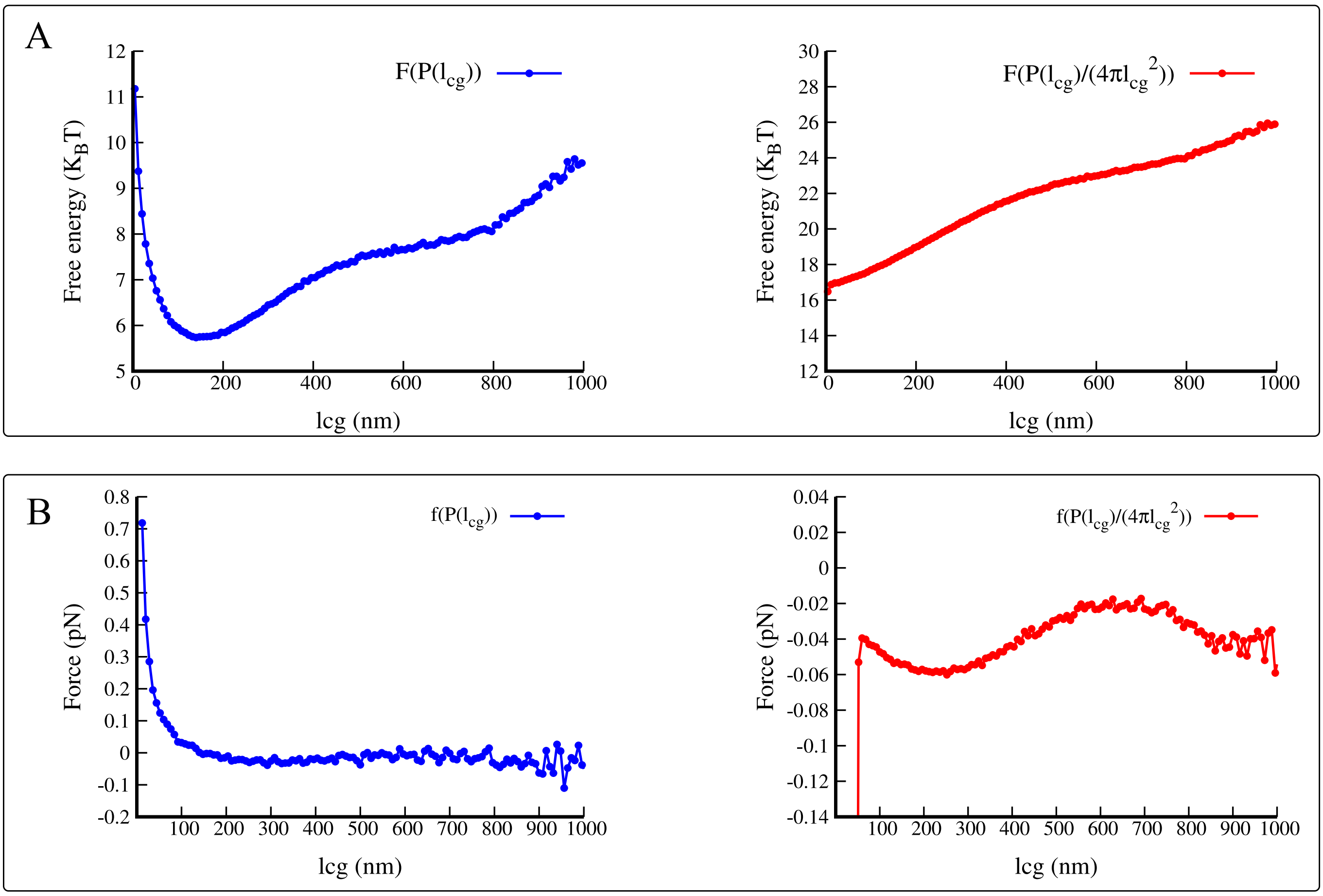


**FIG. S3:** Free energy profiles and corresponding force-extension relations derived from bond-length distributions obtained from microscopy data (Bintu et al. [8]). (A) Free energy as a function of bond length $l_{cg}$ computed from the probability distribution as $F\left(P(l_{cg})\right) = -k_B T \ln P(l_{cg})$ (left). The right panel shows the free energy including the geometric (Jacobian) factor, $F\left(P(l_{cg})/(4\pi l_{cg}^2)\right) = -k_B T \ln\left[P(l_{cg})/(4\pi l_{cg}^2)\right]$. Both are expressed in units of $k_B T$.
(B) Corresponding force-extension relations obtained from the free energy profiles as $f\left(P(l_{cg})\right) = -\frac{d}{dl_{cg}}\left[F\left(P(l_{cg})\right)\right]$ (left) and $f\left(P(l_{cg})/(4\pi l_{cg}^2)\right) = -\frac{d}{dl_{cg}}\left[F\left(P(l_{cg})/(4\pi l_{cg}^2)\right)\right]$ (right), evaluated numerically. Note that the force value at large extension is very small, suggesting that the condensed chromatin seems to unfold (flow). This is very much unlike a spring.

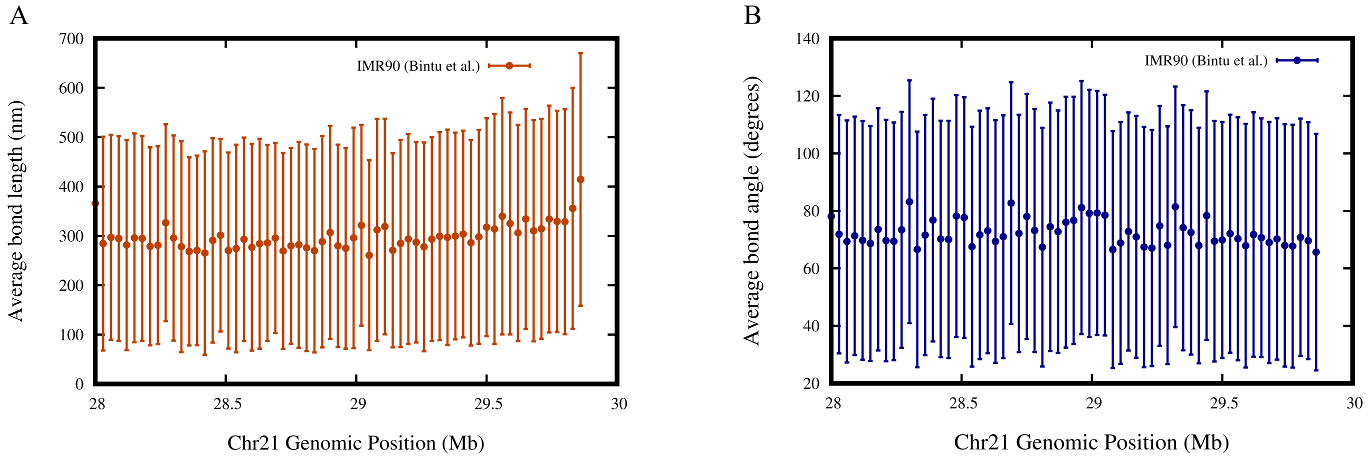


**FIG. S4:** Every chromatin region exhibits high variability in bond length ($l_{cg}$) and bond angle ($\theta_{cg}$) across single cells. (A) Bond length between adjacent 30 kb chromatin segments is plotted along the genomic coordinate (x-axis), with the mean bond length and corresponding standard deviation across single cells shown on the y-axis. (B) Similarly, the mean bond angle and its variability (standard deviation across single cells) are plotted as a function of genomic coordinate. The distributions are computed from microscopy data reported by Bintu et al [8].

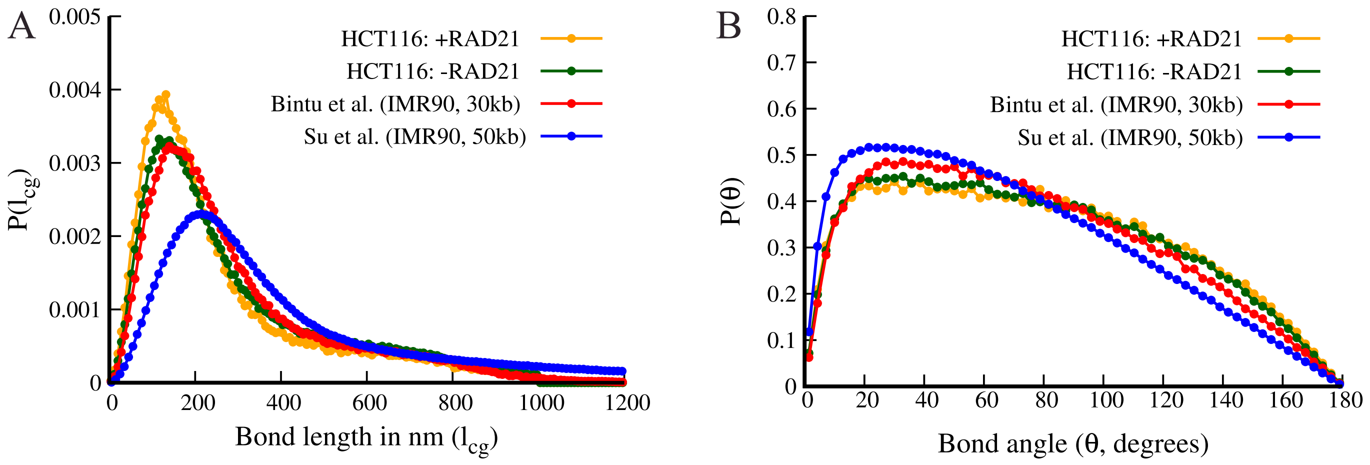


**FIG. S5:** (A) Bond-length distribution and (B) bond-angle distribution derived from microscopy data [8, 11] under different experimental conditions and resolutions. Distributions are shown for the 30 kb resolution in the Chr21: 28-30 Mb region, and for the 50 kb resolution across the entire chromosome 21 [11]. Additionally, data for the HCT116 Chr21: 28-30 Mb region are also plotted under auxin-induced RAD21 depletion (-RAD21) and untreated conditions (+RAD21). This data shows that long tails in bond length distributions and acute angle in angle distribution persist at 50 kb resolution, and are also observed in RAD21-depleted cells.

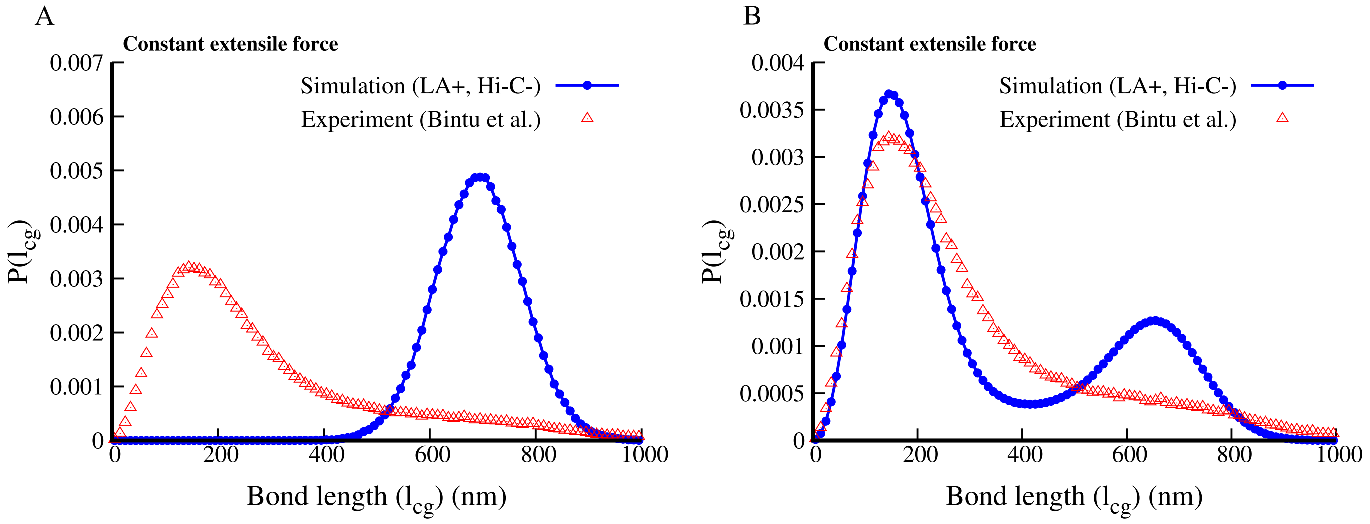


**FIG. S6:** (A) Bond-length distribution obtained from simulations, where a constant extensile force of 5.49 pN is applied uniformly for every neighbouring pair throughout the simulation. (B) Bond-length distribution obtained from ADCM simulations (see methods) in which all parameters are kept identical, except that the extensile force is applied as a constant (5.49 pN) rather than a variable. As described in the ADCM method every triplet switches between extensile and angular forces. This shows that a constant extensile force does not produce a long tail; instead, it either produces a shift or results in a bimodal distribution with two peaks.

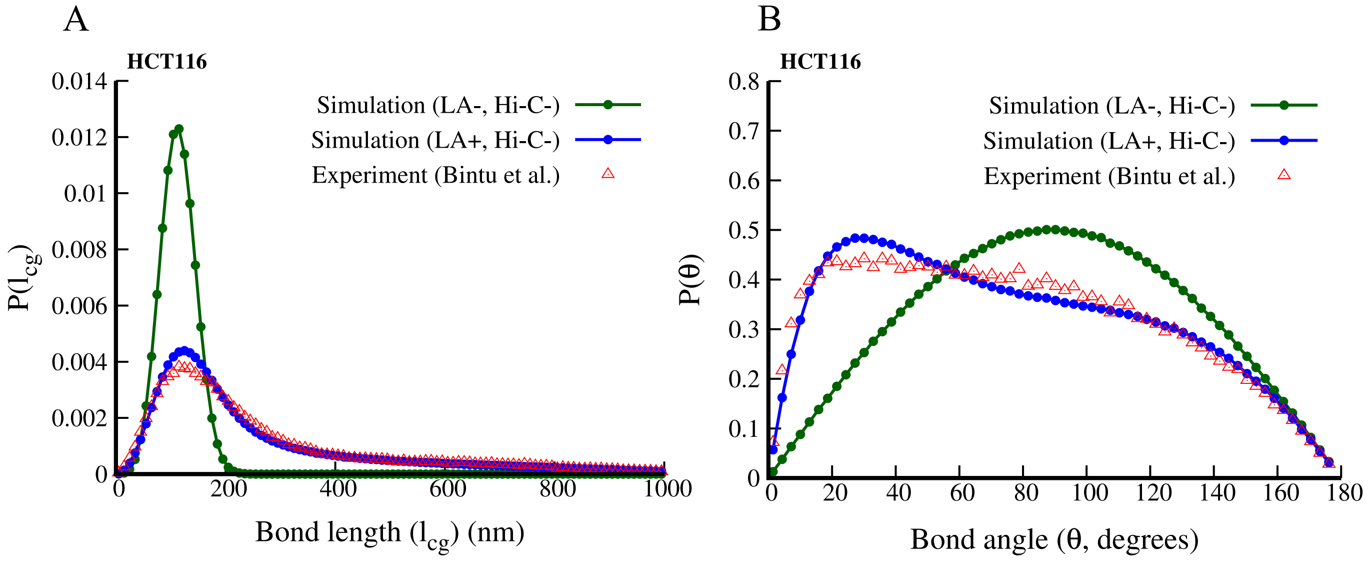


**FIG. S7:** Similar to Fig. 3 in the main text, this figure shows a comparison of bond-length ($l_{cg}$) (A) and bending-angle ($\theta_{cg}$) (B) distributions obtained from simulations and microscopy data [8] for the HCT116 cell line (Chr21: 28–30 Mb region). LA+/LA- denote simulations with and without local active forces, respectively, while Hi-C- indicates that long-range interactions observed in Hi-C (e.g., due to extrusion, CTCF, etc.) are not included.

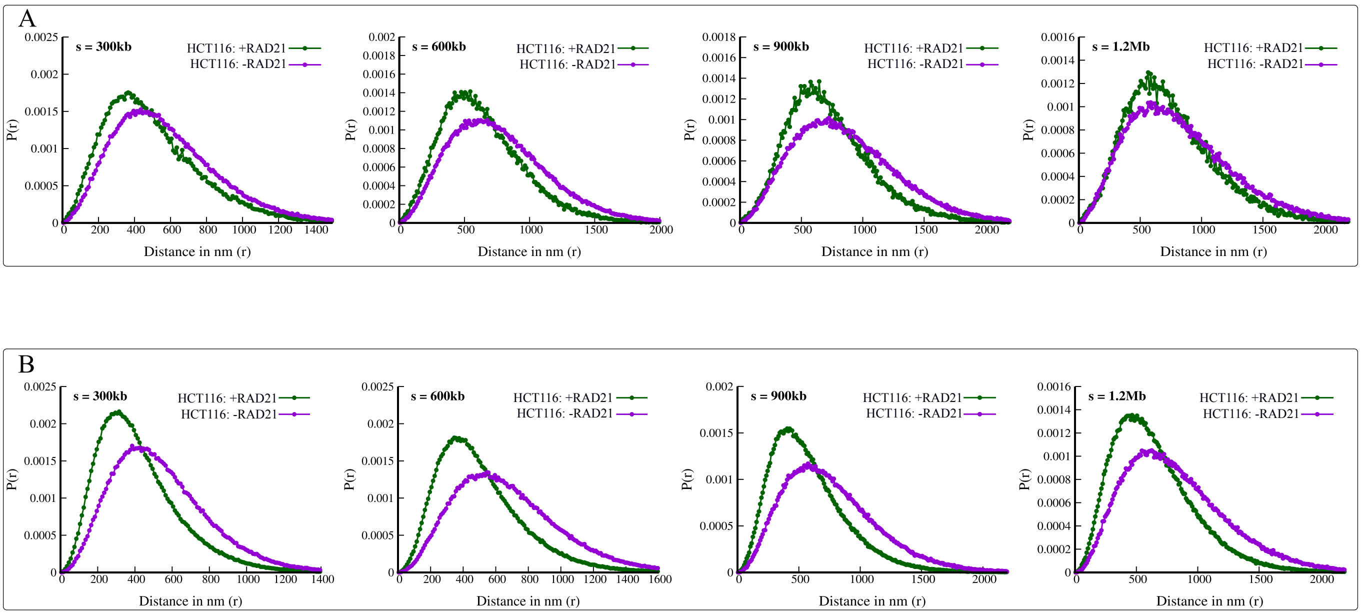


**FIG. S8:** 3D distance distribution P(r) as described in Fig. 3 of the main text. Here, we plot P(r) with and without auxin-induced depletion of RAD21 for (A) the 28-30 Mb region and (B) the 34-37 Mb region of chr21 in the HCT116 cell line, computed from Bintu et al. data [8]. This shows that RAD21 depletion prominently affects 3D distances between any two points that are far away (> a few hundred kb) along the genome.

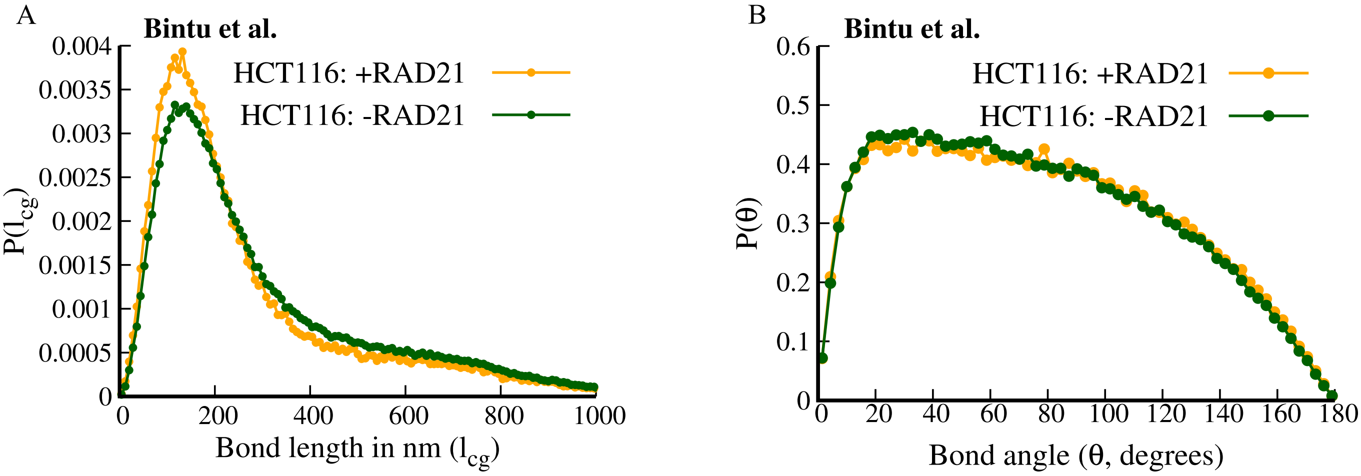


**FIG. S9:** (A) Bond-length (left) and bond-angle (right) distributions for the HCT116 cell line, chr21: 28–30 Mb region, under conditions with auxin-induced depletion of RAD21 (-RAD21) and without auxin treatment (+RAD21), computed from Bintu et al. data [8]. This indicates that loop extrusion does not majorly affect bond-length and bond-angle distribution at 30kb scale.
